# 5′ untranslated regions tune *Toxoplasma* translation

**DOI:** 10.1101/2025.07.14.664749

**Authors:** Michelle L. Peters, Aditi Shukla, Zoe A. Gotthold, Dylan M. McCormick, Kehui Xiang, David P. Bartel, Sebastian Lourido

## Abstract

Some of the longest 5′ untranslated regions (UTRs) documented in eukaryotes belong to parasites of the phylum Apicomplexa. Translational regulation plays prominent roles in the development of these parasites, including the agents of toxoplasmosis (*Toxoplasma gondii*) and malaria. To understand the function of 5′ UTRs in apicomplexan translation, we performed high-resolution ribosome profiling of *T. gondii* in human fibroblasts. We show that parasite translation efficiency (TE) is largely controlled by 5′ UTR features and tuned by the number of upstream AUGs (uAUGs). Examination of ribosome occupancy reveals that, despite widespread assembly of parasite monosomes on uAUGs, ribosomes seldom translate uORFs. These determinants of translation are reaffirmed in a massively parallel reporter assay examining the effect of more than 30,000 synthetic 5′ UTRs in *T. gondii*. A model trained on these results accurately predicted the TE of newly designed 5′ UTRs. Together, this work defines the regulatory language of *T. gondii* translation, providing a framework to understand the evolution of exceptionally long 5′ UTRs in apicomplexans.

## INTRODUCTION

Translation is a tightly-regulated step in eukaryotic gene expression.^1^ A wide range of pathways control translation globally or in a transcript-specific manner, particularly by influencing rates of translation initiation.^2^ In eukaryotes, translation initiation generally occurs through a scanning mechanism in which the preinitiation complex—composed of the small (40S) ribosomal subunit, Met-tRNA, and a set of translation initiation factors—is recruited to the mRNA 5′ cap, where it scans along the transcript until a start codon is recognized and the large (60S) subunit joins to form the 80S ribosome.^3^ Although coding sequence (CDS) and 3′ untranslated region (UTR) elements and poly-A tail length are known to regulate multiple aspects of translation and mRNA homeostasis, 5′ UTRs play a dominant role in translational regulation.

Many 5′ UTR features influence the efficiency of translation initiation, including RNA structures, 5′ UTR length, and composition. In particular, the presence of upstream open reading frames (uORFs) has been shown to strongly affect translation.^4^ uORFs are generally thought to suppress translation of the main downstream open reading frame,^5,6^ and can affect ribosome activity through several distinct mechanisms. In some cases, ribosome activity is modulated by interaction with uORF-encoded peptides like the fungal arginine attenuator peptide,^7^ while other uORFs may waylay ribosomes that fail to bypass the upstream AUG (uAUG) during scanning.^4^ Complex interplay may occur on multi-uORF transcripts such as *ATF4* in metazoans^8^ and *GCN4* in yeast.^9^ For these reasons, uAUGs and the associated uORFs generally occur less frequently than would be expected based on the dinucleotide content of 5′ UTRs of most eukaryotic genomes.^6,10^ Indeed, fewer than 50% of mammalian transcripts and only 15% of yeast transcripts contain any uAUGs.^6,11^

Despite their otherwise compact genomes, some members of the phylum Apicomplexa encode 5′ UTRs with average lengths surpassing those of most other eukaryotes.^12^ Based on transcriptional start site (TSS) mapping, the ubiquitous parasite of warm-blooded animals *Toxoplasma gondii* encodes 5′ UTRs with a median length approximately four times that of human 5′ UTRs (∼800 nt vs. 220 nt).^12^ Unusually extended 5′ UTRs are also found in related malaria-causing parasites, with TSS mapping revealing a median 5′ UTR length of over 400 nt in *Plasmodium falciparum*.^12,13^ These long apicomplexan 5′ UTRs exhibit an unusually high number of uAUGs and uORFs, suggesting little apparent selective pressure against the incorporation of these features into transcripts.^6,12^

Across complex life cycles, these single-celled parasites rapidly adapt to changing pressures between intracellular and extracellular environments, host tissue types, and different host species altogether. This requires dynamic control of protein expression, and translational regulation has been implicated in key life cycle transitions. In both *Toxoplasma* and *Plasmodium* spp., translational repression of stage-specific transcripts is critical for parasite development.^14^ The RNA helicase DOZI silences transcripts in female gametocytes until after fertilization in *P. berghei*,^15^ while the Pumilio-2 RNA-binding protein translationally represses liver stage factors during sporozoite development.^16^ In *Toxoplasma*, the master regulator of chronic bradyzoite differentiation BFD1 is translationally repressed in acute-stage tachyzoites.^17–19^ Eukaryotic translation initiation factors (eIFs) including eIF2α, eIF1, and eIF4E have also been identified as key nodes of translational control for apicomplexans.^19–23^

Despite the importance of translational regulation and the unusual transcript features in these species, few apicomplexan uORFs have been functionally characterized. In *Toxoplasma*, currently the only example of functional uORF-mediated regulation is the arginine transporter *Tg*ApiAT1, which is translationally upregulated in response to arginine scarcity.^24^ To investigate translation globally, ribosome profiling is a powerful technique that involves sequencing ribosome-protected RNA footprints as a readout of ribosome position and translation levels.^25–27^ Ribosome profiling has been applied in *Toxoplasma* studies,^19,23,28,29^ with some efforts to identify translated uORFs. However, prior investigations have focused on the roles of *trans*-acting factors or lack the resolution to discern ribosome interactions with specific *cis*-regulatory mRNA elements. Therefore, we still lack a global framework to explain the tolerance for the unusual 5′ UTR features in parasite transcriptomes.

Here, we survey the landscape of translation in *Toxoplasma* acute stages through high-resolution ribosome footprinting. We define transcript features that determine translation levels and deploy a massively parallel reporter assay to uncover an unexpected mode of regulation by the prevalent uAUGs found in parasite 5′ UTRs.

## RESULTS

### High-resolution ribosome profiling improves gene annotation

High-resolution ribosome profiling requires rapid isolation of ribosome-associated RNA, digestion of unprotected RNA, enrichment of 80S ribosomes, and isolation of protected RNA footprints.^27^ To optimally capture translation in the parasite *Toxoplasma gondii*, we profiled acute-stage tachyzoites from the ME49 reference strain growing in primary human foreskin fibroblasts (HFFs) (**Figure 1A**). Two days post infection, cultures were lysed and flash-frozen for downstream processing of ribosome footprints and total RNA libraries. To avoid bias from poly-A selection of total RNA samples, we performed rRNA-depletion using a combination of custom probes against *Toxoplasma* rRNAs and commercial probes against human rRNAs (**Supplementary Data 1**). Of unique reads mapped to a human and *Toxoplasma* metagenome, an average of 32.6% of total RNA reads and 27.3% of ribosome footprints mapped to *Toxoplasma* across two biological replicates. Even with a type II strain, which maintains a lower burden of infection *in vitro*,^30^ this dataset provides higher relative coverage of the parasite over a previous simultaneous study of type I *Toxoplasma* and host translation (∼15-25% *Toxoplasma* reads).^28^ In addition, our methods aimed to avoid biases that may come from cycloheximide pretreatment^27^ or prolonged harvest during mechanical release of parasites from the host cell, one or both of which have been applied in previous *Toxoplasma* ribosome profiling studies.^19,23,29^

**Figure 1.**
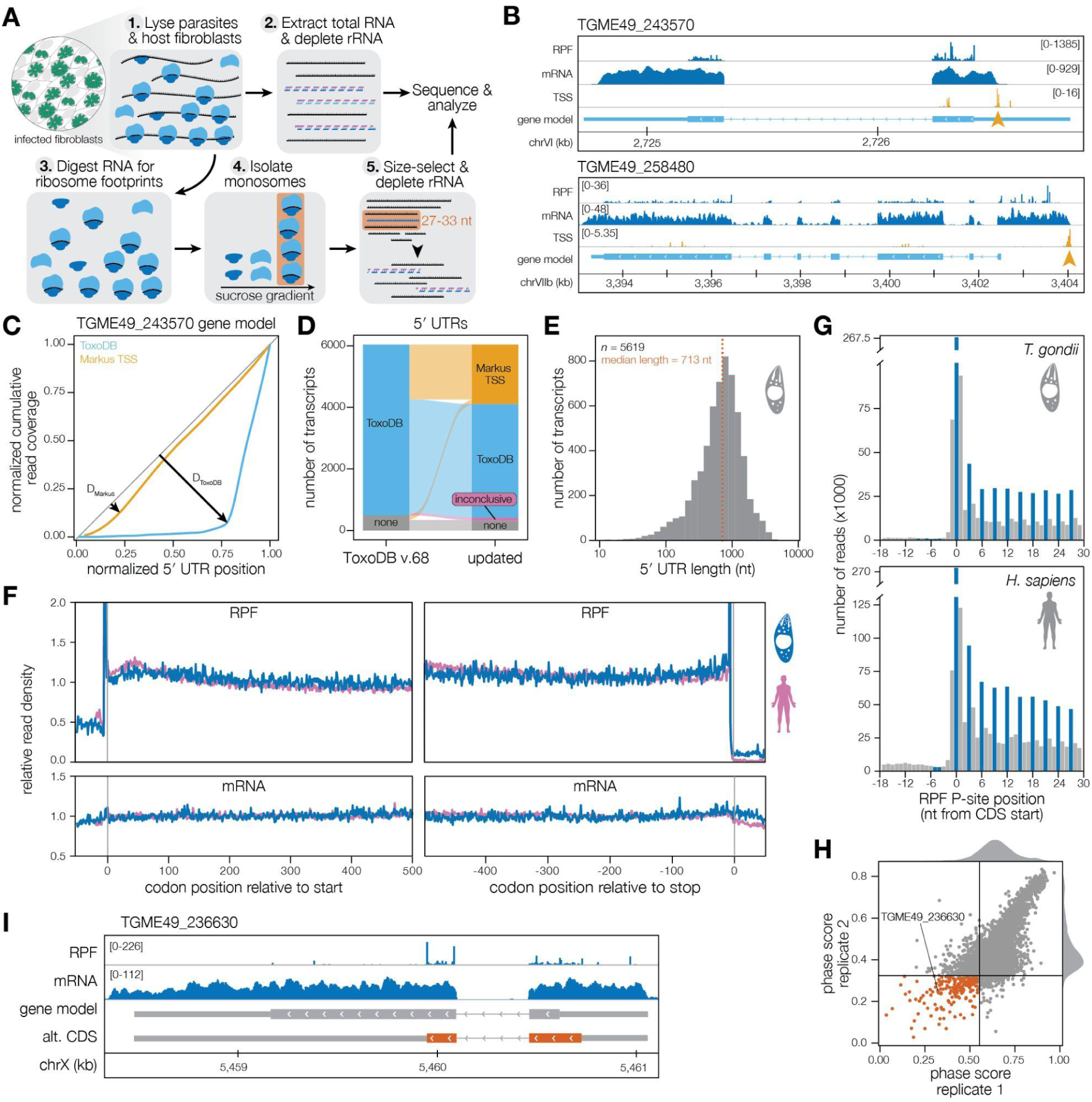
High-resolution ribosome profiling improves *Toxoplasma* genome annotation. (**A**) Library generation of total RNA and ribosome-protected footprints from parasites and host cells. Key fractionation and rRNA depletion steps are illustrated. (**B**) Representative transcripts with discrepancies between ToxoDB gene model and mRNA coverage (top) or lacking a 5′ UTR annotation (bottom). Ribosome-protected fragment (RPF) and mRNA coverage shown in three-nucleotide bins, with a 12-nucleotide correction applied to RPF position. Transcriptional start site (TSS) data from Markus *et al*,^12^ where the orange arrow indicates the called TSS peak. (C) Representative analysis of cumulative read coverage across two 5′ UTR isoforms for *TGME49_243570*. Coverage from total RNA reads was calculated across 3-nucleotide windows of each isoform before normalization for total reads and UTR length. A Kolmogorov–Smirnov test was applied to compare each isoform to an even distribution. The isoform with the smallest differential (D) was selected as representative, except in special cases (see Methods). (D) Sources of 5′ UTR annotations from ToxoDB v. 68 (left) and after curation (right). (**E**) Distribution of *Toxoplasma* 5′ UTR lengths from transcripts with RPKM ≥ 5 after re-annotation. (**F**) Metagene plot of RPFs (top) and total RNA (bottom) for parasite and host. Reads are plotted relative to coding sequence (CDS) start (left) or CDS stop (right). Reads in 3-nt windows were normalized within each gene to the mean coverage per bin, excluding genes with less than 10 reads total. Coverage was then averaged across all genes and across two replicates. (**G**) Nucleotide-resolution plot of reads relative to CDS start after P-site offset correction. In-frame reads are shown in blue, out-of-frame reads are shown in grey. (**H**) Results of ribotricer analysis on *Toxoplasma* annotated CDS. Black lines show phase score cutoffs for translating CDSs for each replicate. Orange points indicate CDSs which failed to pass the phase score cutoff in both replicates. (**I**) Representative transcript identified in **H** with potential alternative CDS.

Examining total mRNA read coverage, we observed discrepancies with many existing gene models, particularly in the annotation of 5′ UTRs (**Figure 1B**). We compared the accuracy of different gene models by drawing on metrics developed to assess mRNA integrity after sequencing.^31^ Incorporating transcript isoforms available in ToxoDB (v. 68)^32^ and transcriptional start sites identified empirically,^12^ we performed a Kolmogorov–Smirnov (KS) test on the cumulative RNAseq read coverage across each possible 5′ UTR for a given gene (**Figure 1C**); the 5′ UTR with the lowest KS statistic (D) was selected as the representative model. These analyses updated 5′ UTR length for 1933 genes, including 152 genes which previously lacked annotated 5′ UTRs (**Figure 1D** and **Supplementary Figure 1A; Supplementary Data 2**). 107 5′ UTRs with high values (D > 0.4) across all models were deemed inconclusive and excluded from downstream analyses. Inconclusive transcripts included lowly expressed genes with poor transcriptional start site signal or those that are incorrectly annotated (e.g. two gene models that should be merged into one); these may require manual annotation or incorporation of additional sequencing data. Of 5619 transcripts with conclusive 5′ UTR models and reads per kilobase per million (RPKM) ≥ 5, we found a median 5′ UTR length of 713 nucleotides (**Figure 1E**). While slightly shorter than prior estimates,^12^ these updated models still place *Toxoplasma* 5′ UTRs amongst the longest observed in eukaryotes, particularly in comparison to humans (median = 220 nt) and model organisms such as *Drosophila* (median = 194 nt) and *Arabidopsis* (median = 185 nt).^12,33^

We next used these updated gene models to map our ribosome profiling data. As expected from previous ribosome footprinting studies across diverse organisms,^26,27,34^ metagene analysis of both parasite and fibroblast transcripts revealed an enrichment of ribosomes in coding regions, with few reads in the 3′ UTRs (**Figure 1F**). After correcting for P-site offsets based on footprint length (**Supplementary Figure 1B**), we observed ribosome protected fragments (RPFs) were enriched at start codons and exhibited trinucleotide periodicity in the CDS, which is consistent with active translation (**Figure 1G**). *Toxoplasma* footprint lengths of 29–30 nucleotides showed the strongest periodicity, and footprints within the range of 27–34 nucleotides were assigned a P-site offset of 12 or 13 nucleotides based on the highest RPF peak accumulating at the CDS start codon for each footprint size (**Supplementary Figure 1B**). Fibroblast RPFs skewed longer than *Toxoplasma* RPFs, which may reflect slightly different requirements for complete RNaseI digestion of unprotected RNA in dual-species ribosome profiling samples.

Given the strong periodicity of *Toxoplasma* RPFs, we wondered whether these data might also aid in confirming CDS annotations in existing *Toxoplasma* gene models. To identify CDSs that may be misannotated, we implemented ribotricer^35^ on annotated coding regions to calculate phase scores. A higher phase score indicates more consistent trinucleotide periodicity across the ORF. We applied phase-score cutoffs based on training for each replicate (**Figure 1H** and **Supplementary Data 3**). Of 4365 transcripts passing a read-density cutoff, 208 had coding regions that failed the phase-score cutoff for both replicates. Although some cases may result from low coverage or overlapping upstream ORFs in a different frame, visual inspection suggests many of these low-phase score genes are incorrectly annotated or have multiple translation start sites. For instance, *TGME49_236630* appears to be translated both in its annotated CDS and, much more strongly, from an alternative ORF (**Figure 1I**). Indeed, the alternative *TGME49_236630* ORF displayed higher homology across the related species *Hammondia hammondi* and *Neospora caninum* (94.24% and 76.81% identity, respectively) than the annotated CDS (89.47% and 61.98% identity, respectively). Furthermore, BLAST searches of the alternative protein sequence against other Apicomplexan genomes identified potential homologues such as *BESB_036630* in *Besnoitia besnoiti*. Although a reannotation of the coding genome is beyond the scope of this manuscript, phase scores from our data can be used to nominate gene models for further evaluation. Taken together, our high-resolution total RNA and ribosome footprint datasets function as a resource to continue improving *Toxoplasma* gene annotations.

### Translation efficiency regulates tachyzoite protein expression

To capture protein translation rates across the *Toxoplasma* and host transcriptomes, we calculated translation efficiency (TE), which is the ratio of RPF RPKM to total mRNA RPKM (**Supplementary Data 4**). TE was calculated from reads aligning to the CDS, excluding the first three and the last three codons to avoid artifacts from translation ramp-up or ramp-down.^27^ While *Toxoplasma* RPF levels generally correlate with mRNA levels (**Figure 2A**), *T. gondii* transcripts exhibit a wide range of translation efficiencies, compared to the narrower range observed for host transcripts (**Figure 2B**).

**Figure 2.**
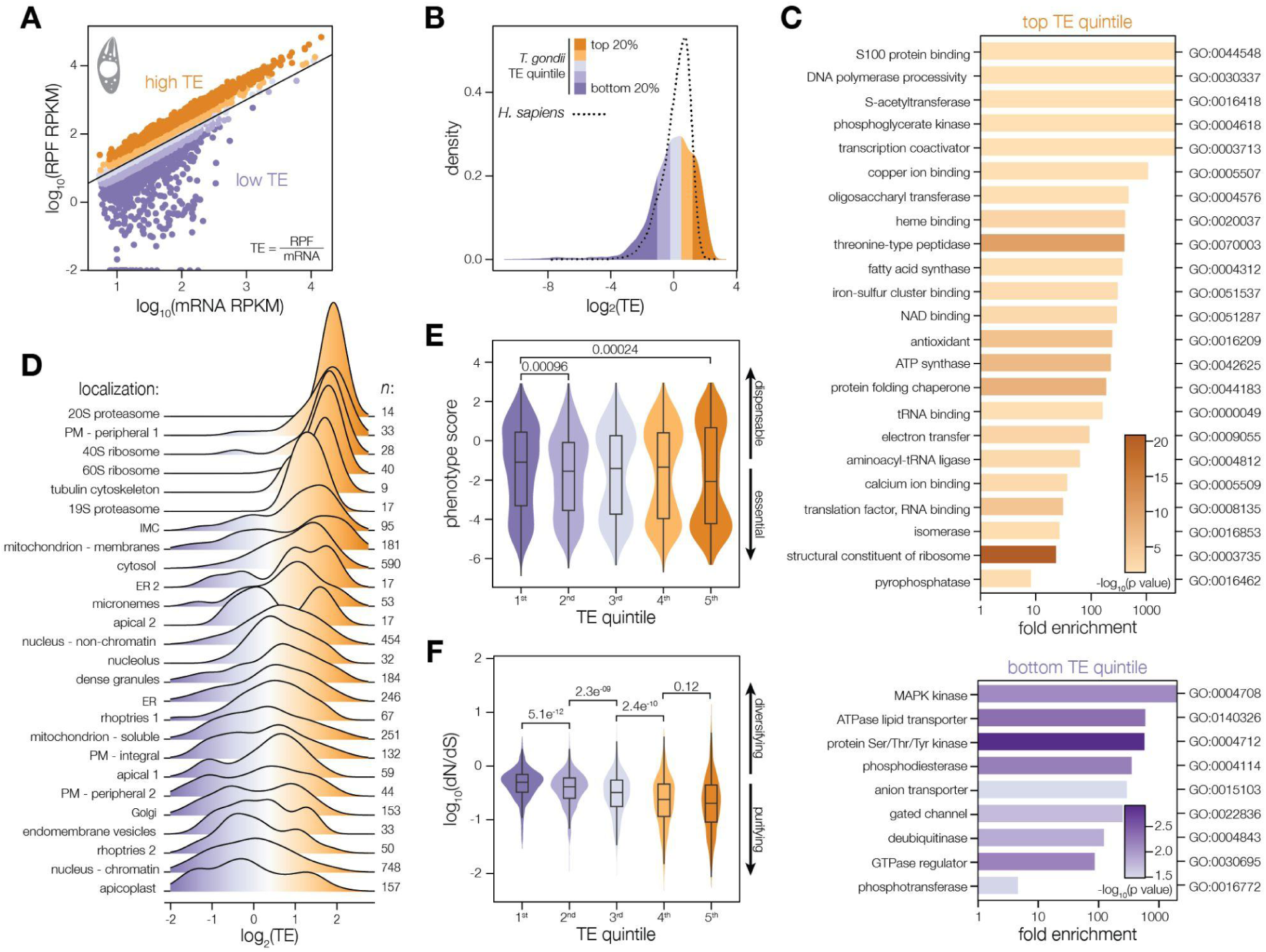
Translation efficiency tunes the composition of the tachyzoite proteome. (**A**) Comparison of ribosome footprint and total RNA levels in the CDS, given in reads per kilobase per million (RPKM). Translation efficiency (TE) is calculated as the ratio of RPF to mRNA reads. Values are the average of two replicates, and solid line represents x=y. Color scale indicates *T. gondii* TE quintiles. (**B**) Distribution of TEs in the parasite and host, with color scale as in **A**. (**C**) Gene ontology (GO) enrichment analysis of top quintile (top) and bottom quintile (bottom) genes as compared to all expressed genes. GO terms were downloaded from ToxoDB, and significance was assessed with a hypergeometric test. Terms with *p* < 0.05 and more than one gene are displayed, with additional manual filtering to remove redundant terms. (**D**) TE distribution of genes in each subcellular localization, as defined by hyperLOPIT TAGM-MAP.^36^ (**E**) Distribution of CRISPR phenotype scores^37^ for genes in each TE quintile. Pairwise Wilcoxon rank sum test with Bonferroni correction. Non-significant (*p*>0.05) comparisons not shown. (**F**) Distribution of dN/dS scores^38^ by TE quintile. Pairwise Wilcoxon rank sum test with Bonferroni correction; all comparisons not shown are also significant (*p* < 0.0001).

We first investigated whether TE is associated with gene function, as represented by gene ontology (GO) terms. Genes from the top or bottom TE quintiles (1199 genes each) were compared to the entire gene set for GO enrichment analysis (**Figure 2C**). The most efficiently translated genes are mainly involved in translation and protein homeostasis, including translation factors, ribosomal proteins, tRNA processing factors, and protein folding chaperones, as well as proteasome components. Respiration-related genes are also enriched in the top TE quintile, including NAD binding proteins, ATP synthase components, electron transfer factors, and heme binding proteins. By contrast, lowly translated genes consist largely of hypothetical proteins (561 out of 1199). Because we did not exclude transcripts with 0 RPF counts from our analysis, some bottom quintile genes may be misannotated or noncoding, resulting in the large population of hypothetical proteins in this group. Nonetheless, genes with low translation efficiency tended to be associated with signaling, such as cyclic nucleotide regulation and kinase activity, as well as transporters and channels. This includes the arginine transporter *Tg*ApiAT1 (*TGME49_215490*), which is translationally activated in response to arginine depletion via a uORF-dependent mechanism.^24^ Similarly, other low TE transporters may be poised for translational upregulation in response to nutrient scarcity.

As an alternative classification of gene functionality, we next analyzed the relationship between TE and protein localization using the hyperLOPIT dataset (**Figure 2D**).^36^ As seen in the GO enrichment analysis, ribosome and proteasome components have amongst the highest TEs on average. Plasma membrane proteins displayed a wide range of TEs. Group PM 1, consisting mainly of surface antigens (SAGs), exhibited high TEs, in contrast to the lower average TEs of PM 2 and integral membrane proteins. ER proteins display a similar pattern, with soluble proteins (ER 2) having higher TEs than integral membrane proteins (ER 1). Thus, TE may coordinate protein levels required across different functional compartments.

Given that highly translated genes are enriched for core cellular functions, we wondered whether TE is correlated with contributions to parasite fitness, as previously measured in a genome-wide CRISPR screen.^37^ Indeed, lowly translated genes have more dispensable phenotypes, while the most highly translated genes are more essential on average (**Figure 2E**). This trend is distinct from that previously reported between total RNA levels and phenotype score, where the most highly expressed RNAs do not have more essential phenotypes on average.^37^ This suggests that TE helps boost levels of proteins that cannot be subject to perturbations in expression without fitness cost to the parasite. On the other end of the spectrum, translational repression may prevent unnecessary production of proteins not required in cell culture or in the tachyzoite stage. For instance, *BFD1*, the master regulator of differentiation to bradyzoites,^17^ is one of the most lowly translated transcripts (bottom 2% TE).

Finally, we evaluated the relationship between TE and selective pressure as measured by the ratio between rates of nonsynonymous (dN) and synonymous (dS) substitutions using previously reported dN/dS scores.^38^ *Toxoplasma* TE was correlated with dN/dS, with more highly translated genes exhibiting stronger purifying selective pressure (**Figure 2F**). Although the average dN/dS score of the most highly translated genes is low, a notable subset—primarily surface antigens and secreted effectors—show signs of diversifying selection. This suggests that despite different evolutionary pressure on coding regions, translational regulation in *Toxoplasma* boosts expression of both conserved functionalities such as protein homeostasis as well as rapidly-evolving parasite proteins like surface antigens.

Collectively, these analyses indicate that transcription and translation work in tandem to shape the tachyzoite proteome. The broad range of TEs in *Toxoplasma* and the significant correlations between TE and functional categories suggest that translation is indeed an important point of regulation determining protein levels in the parasite.

### Different transcript features impact translation efficiency in *Toxoplasma* and human cells

To identify potential *cis*-acting features regulating TE in *Toxoplasma*, we dissected the impact of different transcript regions and compared these between parasites and host cells (**Figure 3A**). Consistent with reports from other eukaryotes,^27,39–42^ we observe a suppression of TE with increasing lengths of 5′ UTR, CDS, or 3′ UTR in our primary human fibroblast samples.

**Figure 3.**
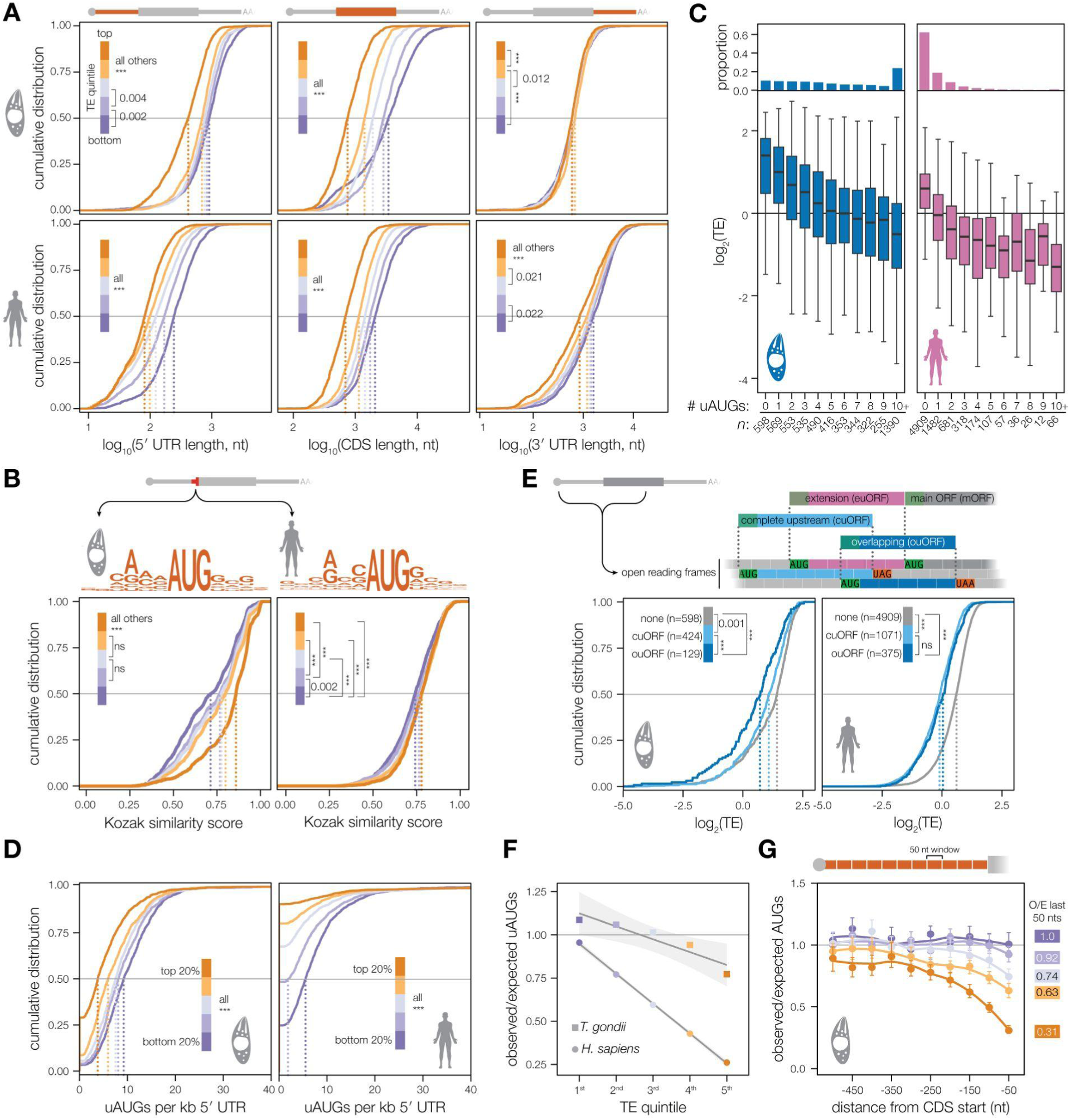
Translation efficiency depends on different sequence elements in *Toxoplasma* and human cells. (**A**) Cumulative distribution of 5′ UTR (left), coding sequence (CDS, middle), and 3′ UTR (right) length, grouped by TE quintile. Pairwise Wilcoxon rank sum test with Bonferroni correction. Non-significant (*p*>0.05) comparisons not indicated. (**B**) Start codon consensus sequences of *T. gondii* and *H. sapiens* (top) above cumulative distribution of Kozak similarity scores across TE quintiles, for each species. Pairwise Wilcoxon rank sum test with Bonferroni correction. (**C**) Bar chart indicates the proportion of the transcriptome belonging to each bin, above the distribution of TEs for transcripts with different numbers of uAUGs. (**D**) Cumulative distribution of number uAUGs normalized for 5′ UTR length. Pairwise Wilcoxon rank sum test with Bonferroni correction. (**E**) The top schematic indicates definitions of ORF types, above the cumulative distribution of TE across transcripts with no uAUGs, one complete uORF (cuORF), or one overlapping uORF (ouORF). Pairwise Wilcoxon rank sum test with Bonferroni correction. (**F**) Observed/expected (O/E) number of uAUGs per 5′ UTR. 5′ UTRs were individually shuffled, maintaining dinucleotide frequency, before calculating overall AUG frequency across transcripts of each TE quintile. Values shown are the median of 1000 simulations. A linear model was fit to O/E values of each species with standard error shown. (**G**) O/E of AUGs across 50 nucleotide windows of the 5′ UTR. 5′ UTR windows were individually shuffled, maintaining dinucleotide frequency. Values shown are the median of 100 simulations with 95% confidence interval.

Intriguingly, *Toxoplasma* maintains a similar relationship only between CDS length and TE, and there is no relationship between 3′ UTR length and TE. The most efficiently translated *Toxoplasma* transcripts generally have shorter 5′ UTRs, but, notably, the distribution of 5′ UTR lengths is nearly indistinguishable among the bottom three TE quintiles. This contrasts with our observations in fibroblasts where we see an inverse correlation between 5′ UTR length and TE across all quintiles. Since length explained only a small portion of TE differences among *Toxoplasma* transcripts, we focused on sequence differences in the 5′ UTRs that may regulate translation.

To assess the impact of start codon context on TE,^43^ we generated a position-weight matrix (PWM) of nucleotides surrounding the translation initiation site (−6 to +4) of all annotated tachyzoite transcripts (**Figure 3B**). This Kozak consensus sequence agrees with previous reports.^12,29,44^ We then calculated a Kozak similarity score for each transcript based on the information content of the PWM.^45^ We found that TE generally correlates with greater Kozak similarity, with the strongest effect observed for the most highly translated transcripts (**Figure 3B**). The same analysis of human transcripts found a small but significant effect for Kozak similarity across all TE quintiles.

Focusing on further 5′ UTR features, we next analyzed the relationship between the number of upstream start codons (uAUGs) and TE. In our updated tachyzoite 5′ UTR models, 90% of *Toxoplasma* transcripts have uAUGs, with a median of 5 uAUGs per transcript (**Supplementary Figure 2A**). By contrast, fewer than 50% of mammalian transcripts contain uAUGs.^46^ The relative scarcity of uAUGs in mammalian 5′ UTRs is generally ascribed to the ability of even one uAUG to strongly suppress translation, which was also evident in our data (**Figure 3C**). By contrast, we observe a gradient of TE suppression across the extreme range of uAUG dosages in *Toxoplasma* (**Figure 3C**). Because the number of uAUGs is strongly correlated with 5′ UTR length in *Toxoplasma*,^12^ we examined whether the relationship between the number of uAUGs and TE is simply a consequence of 5′ UTR length. We still found a significant correlation between the number of uAUGs per kb and TE for both *Toxoplasma* and humans (**Figure 3D**), although the distribution was shifted dramatically higher in *Toxoplasma*. Intriguingly, the most highly translated transcripts in *Toxoplasma* harbored fewer uAUGs per kb. These data suggest that *Toxoplasma* better tolerates uAUGs compared to mammalian cells and that the number of uAUGs tunes TE.

To understand how uAUG context impacts translation, we categorized uORFs based on their relationship to the CDS (designated the main ORF or mORF). We compared three categories: (i) ORFs that start and terminate upstream of the mORF (complete uORFs, or cuORFs), (ii) those that terminate within the mORF (overlapping ORFs, or ouORFs), and (iii) N-terminal extensions, which are in-frame start codons upstream of the annotated CDS (euORFs) (**Figure 3E**). To avoid convoluting the effects of multiple uORFs, we analyzed the subset of transcripts with 0 or 1 uAUGs. EuORFs had no effect on TE in either species (**Supplementary Figure 2B**), which would be expected as ribosomes initiating at these sites could continue to translate the mORF in-frame. Interestingly, ouORFs suppress TE more than cuORFs in *Toxoplasma*, whereas we observe no difference in TE between transcripts with a cuORF and those with an ouORF in human cells. Together, these analyses indicate that the ability of ribosomes to reinitiate following translation of a uORF may be more important in *Toxoplasma* than in humans.

In mammals, uAUGs occur less frequently than would be expected from the dinucleotide frequency of UTRs, suggesting selective pressure against their accumulation.^6^ We compared observed (O) and expected (E) numbers of AUGs in the 5′ UTRs of human and *Toxoplasma* transcripts (**Figure 3F**), deriving the expected numbers from simulations maintaining dinucleotide content.^6^ An O/E ratio less than one indicates selection against uAUGs. Human transcripts in all TE quintiles display evidence of purifying selection, with a strong negative correlation between TE and O/E ratio. By comparison, only the top two *Toxoplasma* quintiles—particularly the top quintile—have a suppressed O/E ratio, and this signal is significantly less pronounced than in humans. Previous studies have shown that selection against AUGs is stronger closer to the CDS start.^47^ Analyzing O/E across 50 nt windows, we indeed observed lower O/E ratios proximal to the CDS in both *Toxoplasma* and humans (**Figure 3G; Supplementary Figure 2C–D**). This effect was increasingly prominent with increased TE; however, the suppression in *Toxoplasma* was limited to a smaller window and effect size. These results extend prior findings,^12^ affirming there is weak, proximity-dependent selection against uAUGs in *Toxoplasma*, and suggesting that uAUG suppression boosts expression of a privileged set of transcripts.

### Ribosomes assemble on upstream AUGs but rarely translate *Toxoplasma* uORFs

Given the impact of uAUGs on TE, we next analyzed how ribosomes engage with uAUGs. Metagene plots of uORF and mORF initiation and termination reveal high ribosome occupancy on uAUGs in *Toxoplasma*, even compared to mAUGs (**Figure 4A**). However, ribosome occupancy following a uAUG peak is much lower relative to the signal within mORFs. To quantify this difference, we calculated an enrichment score of AUG counts over the average counts of the three codons following them. In *Toxoplasma*, the AUG:ORF enrichment score is 23.6x for uAUGs and 7.9x for mAUGs. By contrast, humans have similar ratios of 4.5x and 4.0x for uAUGs and mAUGs, respectively. These observations suggest that while initiation occurs at many *Toxoplasma* uAUGs, these events do not generally progress to elongation on uORFs. Additionally, the proportional ribosome occupancy on start codons is much greater than that on stop codons in *Toxoplasma*, indicating that translation initiation is slower than termination.

**Figure 4.**
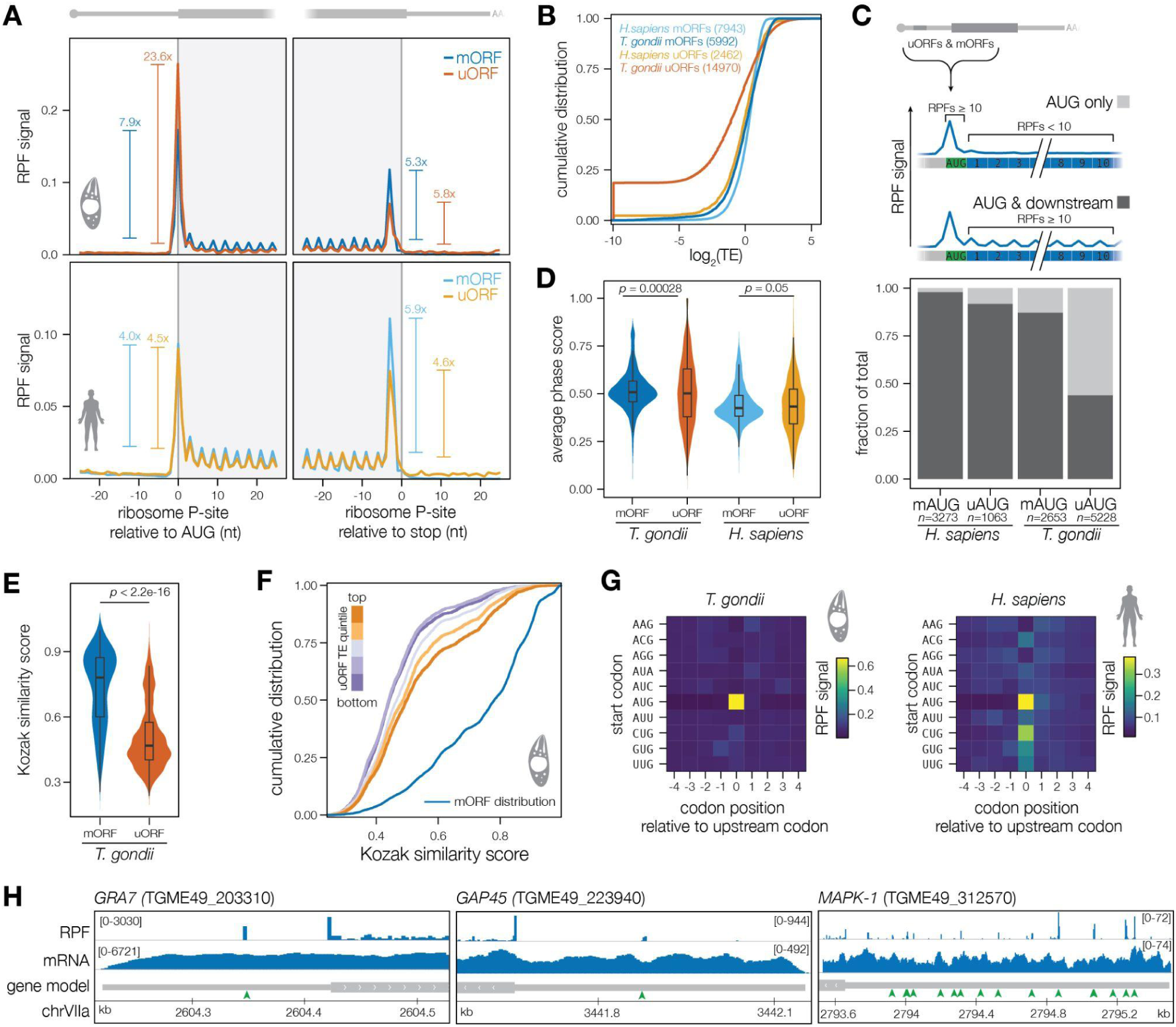
Ribosomes assemble on upstream AUGs but rarely translate *Toxoplasma* uORFs. (**A**) Metagene plots of P-site corrected ribosome protected fragments (RPF) around the start and stop codons of main ORFs (blue) and uORFs (orange). ORFs were included only if there were no other AUG or stop codons within the 50-nt window around the feature. P-site corrected read counts for each position on the uORF or mORF were divided by the count sum for that ORF, before averaging across all features and two replicates. Enrichment values show the ratio of counts on the start (coordinate 0) or stop (coordinate −3) residue divided by the average counts across the first 3 codons (coordinates 3, 6, and 9) or last 3 codons (coordinates −12, −9, and −6) of the ORF. (**B**) Cumulative distribution of translation efficiency for uORFs and mORFs in both species. Only uORFs longer than 45 nucleotides were included, and ouORFs were excluded. TE was calculated as for mORFs, excluding the first 3 and last 3 codons. (**C**) Proportion of uAUGs and mAUGs with ≥10 RPF counts on the AUG and/or within the following 10 codons. RPFs were assigned to the AUG if the P-site fell on nucleotide position −1, 0, or 1 relative to the A of the AUG. RPFs were assigned to downstream codons if the P-site fell on nucleotide positions +2 to +10. (**D**) Average phase scores from ribotricer analysis across two replicates for uORFs and mORFs with >10 read counts. Wilcoxon rank sum test. (**E**) Kozak similarity scores for *Toxoplasma* mAUGs and uAUGs. Wilcoxon rank sum test. (**F**) Cumulative distribution of Kozak similarity scores for uORFs across uORF TE quintiles. mORF Kozak similarity score distribution is shown in blue for reference. (**G**) Metagene plots of P-site corrected RPFs around alternative start codons in the 5′ UTR. For non-AUG codons, windows including AUGs at other positions were excluded from the analysis. RPFs were summarized at the codon level by adding RPF signals from nucleotide positions −1, 0, and 1 for each codon, where 0 is the first nucleotide of the codon, with normalization as in **A**. (**H**) Representative *Toxoplasma* transcripts with ribosome footprint peaks on uAUGs. uAUGs are indicated with green arrows in the gene model.

As an alternative measure of uORF translation, we calculated uORF TE as previously done for mORFs (i.e. excluding the first and last three codons; **Figure 4B**). Consistent with the average ribosome occupancy, *Toxoplasma* uORFs have much lower TE than mORFs, whereas human uORFs and mORFs have similar TE distributions. To further evaluate the distinction between uAUG occupancy and uORF translation, we separately quantified uAUG/uORF and mAUG/mORF occupancy for AUGs that contained at least 10 read counts and belonged to ORFs longer than 10 codons (**Figure 4C**). Downstream codons were considered occupied if the 10 codons immediately following the AUG had a total of at least 10 read counts. By this measure, translational elongation was similar across human mAUGs, human uAUGs, and *Toxoplasma* mAUGs. However, more than 50% of *Toxoplasma* uAUGs that initiate do not show associated downstream read counts. These data are consistent with a model in which *Toxoplasma* ribosomes assemble at uAUGs, but regularly fail to proceed to elongation. We verified that ribosome footprints in the 5′ UTR originate from initiating and translating ribosomes by implementing ribotricer^35^ for the uORFs (**Figure 4D and Supplementary Data 3**). Overall, phase scores were similar between mORFs and uORFs in *Toxoplasma* and human samples. We attribute the modestly lower phase scores in *Toxoplasma* uORFs to uORFs overlapping in different reading frames.

Because ribosomes assemble on both uAUGs and mAUGs but exhibit different patterns at the initiation to elongation transition, we hypothesized that AUG context may distinguish these events. As previously reported,^12,29^ our analysis confirmed that uAUGs in *Toxoplasma* have a much lower Kozak similarity score than mAUGs (**Figure 4E**), as is true in humans (**Supplementary Figure 3A**). *Toxoplasma* uORFs with higher TE also tended to have higher Kozak similarity scores (**Figure 4F**), suggesting that this context may increase the probability of ribosomes elongating after an uAUG.

Other eukaryotes utilize near-cognate start codons to initiate translation.^26,34,48^ We therefore performed metagene analysis with potential alternative start codons, which differ from AUG by one nucleotide (**Figure 4G**). In *Toxoplasma*, RPF signal was low for any start codon other than AUG. By comparison, initiation was evident on alternative start codons, including CUG, GUG, and UUG, in our fibroblast samples, consistent with studies that identified these codons as frequent initiation sites in mouse and human cells.^34,48^ Thus, translation initiation in *Toxoplasma* is highly specific to AUGs.

The specificity and magnitude of ribosome footprints is clearly apparent on uAUGs when examining individual *Toxoplasma* genes (**Figure 4H**). We observed major peaks of ribosome occupancy at uAUGs of genes with shorter 5′ UTRs (e.g., *GRA7* and *GAP45*), as well as those with more complex 5′ UTRs (e.g., *MAPK-1*); however, few of these uORFs showed any RPFs in their coding regions. Interestingly, the height of ribosome peaks tends to be consistent across the 5′ UTR of a transcript, as represented in the *MAPK-1* example. Additionally, as shown across all three representative transcripts, the height of uAUG peaks scales with the magnitude of reads at the mAUG and CDS.

Integrating the pattern of ribosome occupancy across the 5′ UTR with the stepwise suppression of translation by increasing numbers of uAUGs, we propose that *Toxoplasma* uAUGs tune mORF translation by delaying ribosomes on their way to the mORF through a process that generally involves initiation without elongation at uORFs. This implies that *Toxoplasma* may utilize alternative mechanisms at the initiation-to-elongation transition to disassemble the 80S ribosome (the form capable of generating the types of footprints we have analyzed) and resume scanning of the 5′ UTR. This ability to resume scanning is further supported by the consistency of RPF signal across uAUGs and mORFs on individual transcripts. Furthermore, simulations of uORF length reveal that *Toxoplasma* uORFs are nearly as long as would be expected by random sampling based on dinucleotide frequencies (O/E = 1.05, from 100 simulations), whereas humans have much shorter uORFs than expected (O/E = 0.69). Together, these results point specifically to uAUGs, rather than uORFs as a whole, as the major regulatory feature tuning translation in *Toxoplasma*.

### Design of a massively parallel reporter assay to assess regulation by 5′ UTRs

Although ribosome profiling accurately captures translation, identifying sequence features that influence translation is complicated by the sparse sampling of possible sequences by endogenous transcripts. To examine the effect of varying 5′ UTR sequences in a controlled context, we designed a massively parallel reporter assay (MPRA) that combines reporter-based sorting with high-throughput sequencing. MPRAs allow functional characterization of thousands of regulatory elements and have previously been used to study 5′ UTRs in yeast and humans.^49–51^

We designed a fluorescent reporter construct for flow-based sorting of parasites with variable 5′ UTRs. In this case, we used the type I RH strain, which has a much higher transfection efficiency than the type II strain used for ribosome profiling and allows for integration of expanded libraries. Our reporter construct includes two expression cassettes (**Figure 5A**). The first cassette encodes a constitutively expressed red-fluorescent protein (mKate2) that is separated via a slip peptide (T2A) from the pyrimethamine-resistant allele of *DHFR*, (mKate2-T2A-DHFR), allowing for drug selection and gating on parasites that integrate the reporter. The second cassette encodes monomeric Neon Green (mNG) driven by the constitutive *TUB1* promoter followed by tandem restriction sites that allow for Gibson assembly of a 5′ UTR library. The reporter construct also includes homology arms to a neutral locus on chromosome VI for targeted integration into the genome.^52^ Although homologous recombination is likely less efficient than random integration, it controls for the influence of chromosomal context. We found that uniform mNG expression required the first 50 nt of the *TUB1* 5′ UTR (**Supplementary Fig. 4A-B**). Nonetheless, integration of a small set of 5′ UTRs into reporter constructs generated a range of mNG expression levels that was correlated with the TE of the endogenous 5′ UTR (**Supplementary Fig. 4A-B**).

**Figure 5.**
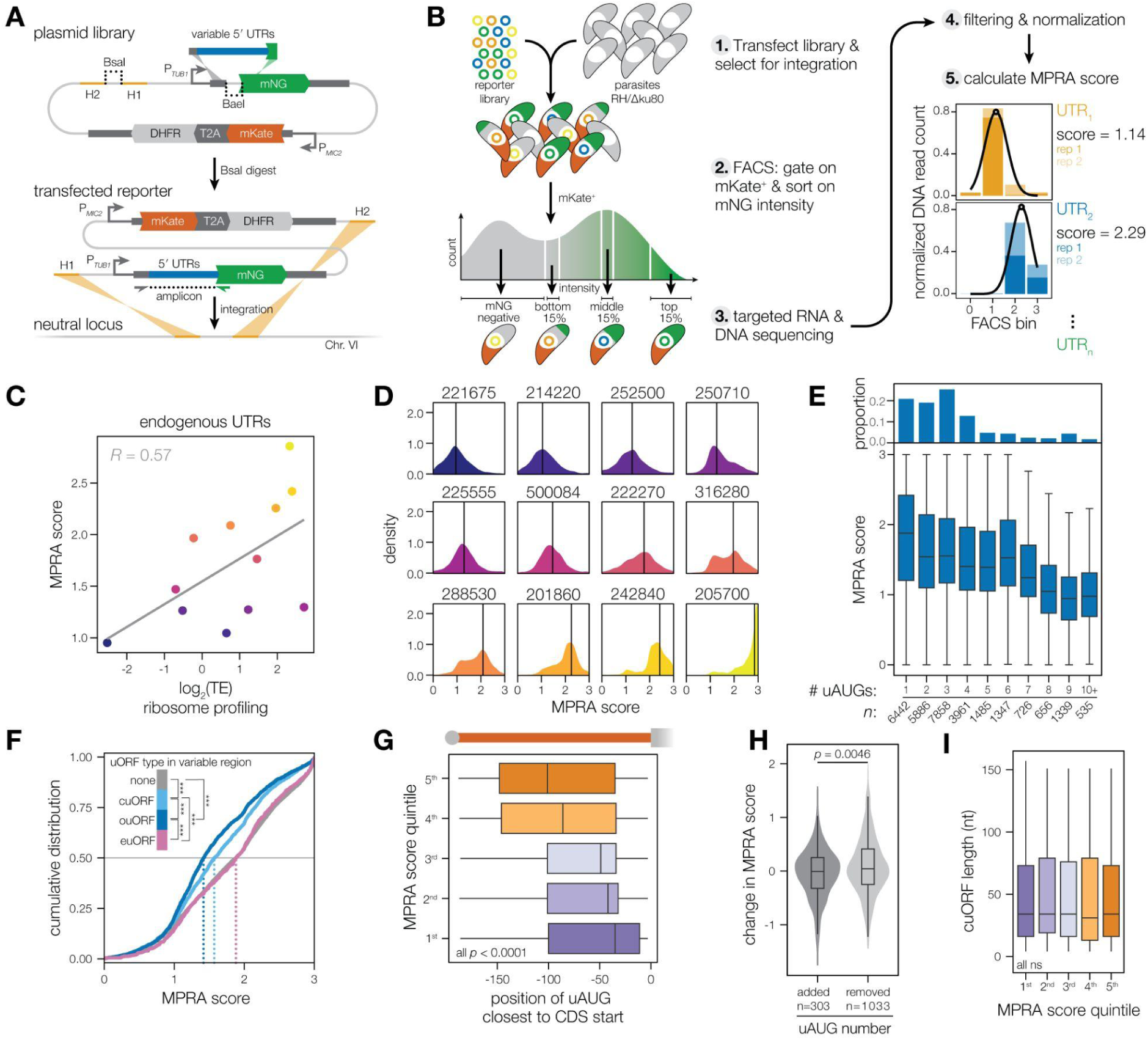
A massively parallel reporter assay (MPRA) reaffirms the regulatory effect of uAUGs on *Toxoplasma* translation. (**A**) Schematic of plasmid for library assembly (top) and linearized reporter with homology arms for chromosomal integration (H1 and H2, bottom). Locations of primers for amplicon sequencing are indicated. (**B**) Experimental and analysis workflow for MPRA showing two representative 5′ UTRs. (**C**) Comparison of MPRA score with TE as measured by ribosome profiling for unmutated 5′ UTRs included in the library. Pearson correlation is shown. (**D**) Distributions of MPRA scores for 5′ UTRs in clades assigned based on similarity to the endogenous UTR. Sequence similarity was calculated with *stringdist* using Levenshtein distance, and sequences with a distance ≤20 from the endogenous UTR were assigned to that clade. The solid line indicates the unmutated UTR’s MPRA score from **C**. (**E**) Cumulative distribution of MPRA scores for 5′ UTRs with different numbers of uAUGs. THe bar chart (top) indicates proportion of the library belonging to each bin. (**F**) Cumulative distribution of MPRA scores for 5′ UTRs with 0 or one uAUG in the variable region (i.e. excluding the constant *TUB1* region), categorized by uORF type. Pairwise Wilcoxon rank sum test with Bonferroni correction. Comparisons not shown are non-significant (*p*>0.05). (**G**) Location of closest uAUG to CDS start, where 0 is the CDS start and negative values are further to the 5′ end of the UTR, by MPRA score quintile. Pairwise Wilcoxon rank sum test with Bonferroni correction. (**H**) Comparison of effects on MPRA score between pairs of sequences with only a change in uAUG number. UTRs with one uAUG in the variable region were used as queries to identify corresponding sequences with point mutations resulting in one additional uAUG, or deletion of the original uAUG. Wilcoxon rank sum test. (**I**) Length of complete uORF (cuORF) in 5′ UTRs with only one cuORF in the variable region, excluding sequences with other types of uORFs. Pairwise Wilcoxon rank sum test with Bonferroni correction.

To design the library of 5′ UTRs, we selected 12 endogenous UTRs covering a wide range of TEs and features (such as uAUGs) that fell within the length limitations of massively parallel oligo synthesis (170–190 nucleotides). For each endogenous UTR, we designed a range of mutations that added or removed uAUGs, entire uORFs, or stop codons; deleted, shuffled, or randomized sequence windows across the UTR; or combined regions of different 5′ UTRs. This resulted in a library of 50,000 sequences that were cloned into the reporter construct.

The assembled library was transfected into a large population of parasites. Over two consecutive passages following drug selection, parasites with different levels of reporter expression were isolated by fluorescence-activated cell sorting (FACS). Parasites gated on mKate expression were sorted into four bins: mNG-negative parasites (1) or mNG-positive parasites in the bottom 15% (2), middle 15% (3), or top 15% (4) of the distribution (**Figure 5B**). After targeted sequencing, we performed quality control and processing, including removing 5′ UTRs with significant discrepancies between bulk DNA and RNA levels to exclude effects originating from RNA stability rather than translation. DNA read counts of each unique 5′ UTR were then used to model a distribution of parasites across the four bins to calculate an MPRA score that represents the average mNG intensity of parasites transfected with that 5′ UTR. We assigned MPRA scores to 30,235 unique 5′ UTRs from the library, allowing us to examine relationships between these diverse sequences and mNG translation (**Supplementary Data 5**).

### MPRA reaffirms the impact of uAUGs on *Toxoplasma* translation

We examined how MPRA scores relate to endogenous TE levels. As expected, the MPRA scores of the 12 endogenous UTRs were correlated with the TEs obtained from ribosome profiling (**Figure 5C**), indicating that the 5′ UTRs recapitulate a significant portion of endogenous expression. We also looked at trends within clades of library sequences that were up to 20 nt different from the endogenous UTR they were derived from. The 12 clades displayed distinct patterns of reporter expression and comprised distributions of MPRA scores that centered around their endogenous UTR of origin (**Figure 5D**). These results indicate that most mutations do not have a large effect on expression; however, certain mutations successfully perturbed native expression levels for each endogenous UTR, suggesting that pathways for modulating TE are robust to sequence context.

We next investigated whether previously described endogenous 5′ UTR features had similar effects in the MPRA. Akin to our TE analysis, we compared the distribution of MPRA scores between subsets of sequences with different numbers of uAUGs (**Figure 5E**). All library constructs contained at least one uAUG from the small portion of the *TUB1* 5′ UTR included in every sequence. As seen in the endogenous context, increasing the number of uAUGs resulted in a stepwise suppression of the MPRA score. Similarly, ouORFs in the MPRA were more suppressive than cuORFs (**Figure 5F**). Furthermore, we observed that reporters with higher MPRA scores tended to have uAUGs further away from the CDS start (**Figure 5G**), in agreement with the apparent suppression of uAUGs towards the start of the CDS in high TE transcripts. We next compared the effect of either adding or removing an uAUG in 5′ UTRs with one uAUG in the variable region and observed that adding an uAUG significantly decreased MPRA scores compared to the removal of a uAUG (**Figure 5H**). By contrast to the significance of uAUGs, we observed no relationship between uORF length and MPRA score in 5′ UTRs with one cuORF in the variable region (**Figure 5I**). Finally, Kozak context was also associated with increased expression of MPRA constructs with motif discovery by STREME^53^ showing an enrichment for optimal Kozak sequences among the top quintile of MPRA-scoring sequences compared to the bottom quintile (**Supplementary Figure 4C**). These results demonstrate a broad consensus between 5′ UTR features that influence TE in endogenous and synthetic sequences.

### A machine learning model trained on MPRA results accurately predicts translation efficiency

To systematically consider a larger number of potential 5′ regulatory features, we trained a random forest model on the MPRA sequences and scores. Adapting methods previously developed for 5′ UTR engineering in human cells,^50^ we scored a range of features including number of uAUGs and uORFs, uORF length, uAUG distance from CDS start, uORF codon usage, RNA folding energy, Kozak score, and K-mers of 1 to 6 nucleotides. For comparison, we also trained a separate model by extracting the same features from endogenous 5′ UTRs and predicting ribosome profiling TE. Cross-validation was performed by training each model using 80% of the data and examining the predictions for the 20% of data withheld over each round of training. Median Spearman correlation values between actual and predicted scores were 0.51 for the ribosome profiling model and 0.66 for the MPRA model (**Supplementary Figure 5A and Supplementary Data 6**), demonstrating that isolating 5′ UTRs in the MPRA indeed allows for better prediction of feature importance.

Both the ribosome profiling and MPRA models emphasize the importance of previously identified features including number of AUGs, AUG positioning, Kozak context, and other uORF-related features (**Supplementary Data 6**). For further insight into the MPRA model, we implemented *fastshap*^54^ to compute approximate Shapley values, which describe the contribution of individual features to a prediction by the model. Globally, top predictors were related to nucleotide content; in particular predicted MPRA scores were positively correlated with U-richness and negatively correlated with G-richness (**Figure 6A and Supplementary Figure 5B**). Multiple measures of RNA folding energy also contribute significantly to the MPRA model, agreeing with a prior report of correlation between translational regulation and RNA structure in *Toxoplasma*.^29^ Overall, the MPRA-trained model reaffirms the importance of previously identified regulatory features, most notably the number and position of uAUGs. Moreover, these data point to new features such as nucleotide content that may play underappreciated roles in regulating translation efficiency in *Toxoplasma*.

**Figure 6.**
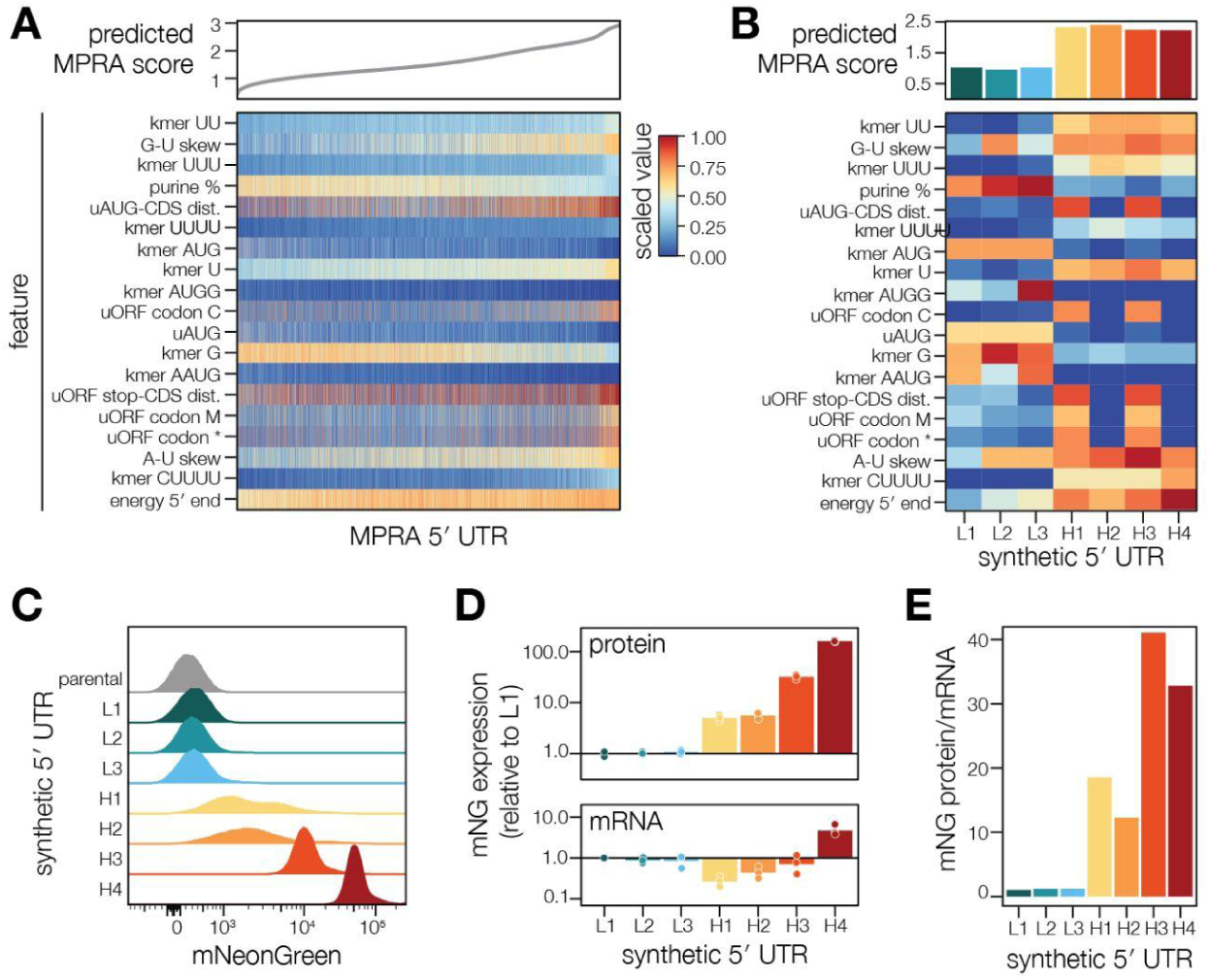
A machine learning model trained on MPRA results accurately predicts translation efficiency. (**A**) Rank order plot of predicted MPRA score by UTR (top) with heatmap of feature values for each UTR (bottom). The top features by average absolute Shapley value are shown. Feature values were scaled from 0 to 1 within each category. (**B**) Features of new 5′ UTRs designed through the genetic algorithm, searching for low-TE (L1-3) or high-TE (H1-3) UTRs. Top barchart displays the MPRA score predicted by the model. Feature values are scaled as in **A** relative to the original MPRA library. (**C**) Representative mNG intensity of UTR reporters as measured by flow cytometry. mNG signal for mKate-positive parasites is shown for all samples except the parental strain, which serves as a background control. (**D**) Quantification of mNG protein levels measured by flow cytometry and mRNA levels measured by RT-qPCR. Points represent biological replicates. RT-qPCR analysis was performed using the 2^−ΔΔCt^ method with *ACT1* as the housekeeping target and L1 as the reference sample. Flow cytometry measurements (median fluorescence intensity) were also normalized to L1 levels. Raw C_t_ values are provided in **Supplementary Data 7**. (**E**) Ratio of protein/mRNA levels for each construct from values in **D**, representing a pseudo-TE score.

To test the predictions of our MPRA-trained model, we used a genetic algorithm (GA)^55^ to generate synthetic UTRs predicted to have low or high TE.^50^ As starting points for the GA, we generated random sequences of 180 nucleotides or randomly selected 180 nucleotide windows of endogenous 5′ UTRs. Following the GA simulations, we curated seven 5′ UTRs from among the highest and lowest predicted TEs to synthesize and test using our reporter construct. We refer to these sequences as L1–3 for predicted low-TE UTRs and H1–4 for predicted high-TE UTRs (**Figure 6B**).

Following transfection and selection, parasites expressing each of the seven expression constructs were analyzed by flow cytometry and reverse transcription and quantitative PCR (RT-qPCR). As expected, expression of the control fluorophore mKate was similar across all constructs at the mRNA and protein level (**Supplementary Figure 5C–D**). By contrast, a clear range of mNG expression levels was observed across the different constructs. The predicted low-TE constructs displayed mNG levels comparable to background from the non-fluorescent parental strain (**Figure 6C**). However, all four predicted high-TE constructs had measurable mNG intensity, with constructs H3 and H4 exhibiting particularly strong expression levels. Critically, these effects originate from differences in translation because all seven constructs expressed the mNG transcript at comparable levels as measured by RT-qPCR (**Figure 6D and Supplementary Data 7**). The greatest variability in mRNA levels was found among the high-TE constructs, with H1 and H4 showing between half and 5 times the mRNA levels in L1, respectively. Taking mRNA levels into account, translation levels of the four high-TE constructs are ∼10 to 40–fold higher than the low-TE constructs (**Figure 6E**). Together, these data demonstrate the mRNA features learned in our model indeed capture regulatory information that can be generalized to new contexts in *Toxoplasma* translation.

## DISCUSSION

Although translational regulation is critical for parasite development and transmission, a global picture of apicomplexan translation has been lacking. Using ribosome profiling and bioinformatic analyses, we provide the first high-resolution global analysis of translation in *Toxoplasma*. We find that *Toxoplasma* transcripts display a wide range of translation efficiencies that are largely driven by *cis*-regulatory features in the 5′ UTR. Distinct from mammalian regulatory regimes, *Toxoplasma* exhibits a gradient of translational suppression in response to uAUG dosage. In line with this correlation, we show that ribosomes assemble on uAUGs but regularly fail to translate uORFs. Through a massively parallel reporter assay, we reinforce the role of uAUGs in setting protein expression levels and identify additional 5′ UTR regulatory elements. Together, our data support a model wherein uAUGs tune translation efficiency by delaying ribosomes during scanning, thereby reducing the rate of translation initiation.

In yeast and mammals, uORFs often function as regulatory elements that license translation of the mORF in response to specific stimuli during development or stress responses.^56^ These mechanisms usually operate on specific transcripts. In *Toxoplasma*, we observe a small subset of uORFs are similarly translated, including the previously-characterized *Tg*ApiAT1 uORF, which regulates expression of the arginine transporter in response to amino acid availability.^24^ However, the extraordinary prevalence of uORFs in *Toxoplasma*—present in 90% of transcripts—necessitates a global framework for their role in parasite translation. Our results corroborate a prior observation that uORFs are lowly translated,^29^ adding codon-level resolution to demonstrate that uAUGs are still occupied by ribosomes despite inefficient uORF translation. These results suggest that the majority of uAUGs in *Toxoplasma* do not function in a stimulus-responsive manner and rather serve to regulate the basal translation levels of mORFs. Although Kozak context partially explains which uAUGs are favorable for elongation and characterizes efficiently translated main ORFs, additional sequence features must distinguish the uAUGs that are initially bound by ribosomes. Nevertheless, the stepwise suppression of translation by increasing numbers of uAUGs argues for a central regulatory function for these features that tunes *Toxoplasma* protein expression.

Further exploration is required to identify the proteins that interact with these mRNA features to regulate ribosomal transitions between scanning, initiation, and elongation amidst the plethora of AUGs. Prior studies of eukaryotic initiation factors (eIFs) in *Toxoplasma*, including eIF4A, eIF4E, eIF2α and eIF1.2, have focused on their roles in differentiation between the acute tachyzoite and chronic bradyzoite stages.^19,22,23,57^ Consistent with the role of eIF1 orthologs in other eukaryotes,^3^ eIF1.2 appears to regulate start codon selection by the *Toxoplasma* PIC,^19^ though the function of its tachyzoite paralog eIF1.1 is yet to be investigated, Curiously, the nearly identical sequences of eIF1.1 and eIF1.2 do not immediately imply any functional distinction. In contrast to other eukaryotes, our data suggest that start codon selection is highly specific to AUGs rather than near-cognate start codons; it may be an adaptation of eIF1 or other factors such as eIF5 and eIF1A that confers this translation start site specificity.^58,59^ More detailed biochemical investigation of *Toxoplasma* ribosomes and initiation factors may clarify the contexts that influence 80S assembly on particular AUGs and license translation elongation almost exclusively on mORFs.

Our study raises further questions on the initiation-to-elongation transition in *Toxoplasma*. Several models could explain the enrichment of ribosome footprints on uAUGs and lack of footprints in uORFs. First, the peaks may reflect a long ribosome dwell time on uAUGs and relatively swift translation of the proceeding uORF. However, given the read coverage in *Toxoplasma* mORFs and human uORFs and mORFs, such a model would require either an extraordinarily slow transition to elongation for *Toxoplasma* uORFs or faster elongation on uORFs than that observed on mORFs. Second, monosomes could assemble on uAUGs but disassemble without proceeding to elongation. Disassembled ribosomes may either fully dissociate from the transcript or permit the small subunit to continue scanning. Based on the consistency of ribosome footprint magnitudes at uAUGs across a 5′ UTR and in the CDS, our data suggest that *Toxoplasma* ribosomes have a high propensity to continue scanning after uAUGs. As with start codon selection, detailed studies of re-initiation are required to confirm this model and identify the proteins that regulate monosome disassembly and continued scanning. For instance, modulation of eIF5B GTPase activity could regulate the fate of ribosomes at this transition in a manner that is responsive to the Kozak sequence.^60^

In addition to *Toxoplasma*, other apicomplexan parasites such as *Plasmodium* spp. maintain extreme numbers of uAUGs in their 5′ UTRs. Similar to our findings in *Toxoplasma*, a study using a luciferase reporter-based system in *P. falciparum* suggested that translation occurs by a combination of leaky scanning, which is dependent on the uORF Kozak sequence, and reinitiation after translation of the uORF.^61^ Polysome profiling^62^ and ribosome profiling studies^63^ of *P. falciparum* revealed high ribosome occupancy in 5′ UTRs. Although the ribosome profiling study argued that ribosome footprints in the 5′ UTR were not specifically enriched in uORFs, it is unclear whether this could be attributed to initiation at alternative codons or artifacts from sample preparation methods, as no periodicity metrics are provided. Nonetheless, 5′ UTR ribosome density decreased in translationally down-regulated genes and increased in translationally up-regulated genes across life cycle stages,^63^ analogously to our finding that uAUG ribosome occupancy scales with mAUG and CDS occupancy. Thus, both *Plasmodium* and *Toxoplasma* may share a common regulatory framework in which translation levels are set by ribosome loading and tuned by sequence elements that impact processivity through the 5′ UTR. Understanding mechanisms of translational control conserved across these parasites but distinct from those of mammals could provide leads for drug development.

Through a massively parallel reporter assay (MPRA), we verified the high tolerance of *Toxoplasma* for uAUGs and their additive effect on translation. To our knowledge, this is the first implementation of an MPRA in an apicomplexan parasite. We demonstrated the feasibility of this powerful technique, which can be readily applied to future studies of *Toxoplasma* gene expression. One limitation of our approach is the length constraint of massively parallel oligo synthesis, which is shorter than the median *Toxoplasma* 5′ UTR length. Future studies could implement alternative library construction strategies to generate longer variable regions. Despite this caveat, leveraging our high-throughput assay and a machine learning pipeline revealed additional features with a strong impact on translation efficiency, which we verified through the generation of novel 5′ UTRs. Most notably, our model and synthetic UTRs demonstrated that in addition to uAUGs, U and G nucleotide content are important for *Toxoplasma* translation. Intriguingly, several recent studies also identified U and G nucleotides as having positive and negative effects, respectively, on TE in other eukaryotes including humans, mice, and zebrafish,^64,65^ while another study of human translation initiation identified C-rich sequences as repressive and AU-rich sequences as enhancing.^66^ An *in vitro* study of *Plasmodium* translation similarly identified a suppressive effect by GC content.^67^ Certain motifs may recruit RNA-binding proteins that are broadly conserved in eukaryotes or alter the biophysical properties of transcripts, which could be especially important in *Toxoplasma* given the length of its 5′ UTRs. Additionally, GC-rich regions might form more stable secondary structures that impede scanning.^68^ Finally, *Toxoplasma* 5′ UTRs were reported to have relatively high levels of pseudouridylation,^69^ which may confer further effects on mRNA stability and translation.

In summary, our work demonstrates that *Toxoplasma* has adapted an unusual mode of translational regulation that largely relies on uAUGs to tune basal translation rates. The extended 5′ UTRs of *Toxoplasma* and related apicomplexans may therefore provide a space for tuning gene expression on a gene-by-gene basis in a more efficient and precise way than could be achieved through changes in promoter elements. This model opens new avenues for work to understand the molecular mechanisms governing ribosome activity in these parasites.

## MATERIALS & METHODS

### Key resources table

| Reagent type (species) or resource | Designation | Source or reference | Identifiers | Additional information |
| --- | --- | --- | --- | --- |
| Strain, strain background (*T. gondii*) | ME49 | PMID:<br>**37582867** | | ME49/*TIR1*/$\Delta$<br>*ku80*/Δ*hxgprt* |
| Strain, strain background (*T. gondii*) | RH | PMID:<br>**19218426** | | RH/Δ*ku80*/Δ*h*<br>*xgprt* |

### Parasite and host cell culture

*T. gondii* were grown in human foreskin fibroblasts (HFFs) at 37°C and 5% CO_2_. ME49 parasites were maintained in DMEM (GIBCO) supplemented with 2 mM L-glutamine (GIBCO), 10 ug mL^-1^ gentamicin (Thermo Fisher), and either 3% or 10% heat-inactivated fetal bovine serum (FBS) where noted (referred to as D3F or D10F). RH parasites were maintained in media supplemented with 2 mM L-glutamine (GIBCO), 10 ug mL^-1^ gentamicin (Thermo Fisher), and 3% inactivated fetal calf serum (IFS) (referred to as D3C) except during transfections and drug selection, for which D10F was used. For routine passaging of ME49 and harvests of intracellular *Toxoplasma*, parasites were mechanically released from host cells by scraping and passage through a 27-gauge needle.

### *T. gondii* transfection

If still intracellular, parasites were mechanically released from host cells. Parasites were then passed through a 5 uM filter to remove host cell debris and pelleted at 1000 x *g* for 5-10 minutes and resuspended along with DNA in Cytomix (10 mM KPO4, 120 mM KCl, 150 µM CaCl2, 5 mM MgCl2, 25 mM HEPES and 2 mM EDTA) supplemented with 2 mM ATP and 5 mM glutathione to a final volume of 400 uL (routine transfections) or 600 uL (large-volume transfections for MPRA). Electroporation was performed with 4-mm cuvettes and an ECM 830 Square Wave electroporator (BTX) with settings: two pulses, 1.7 kV, 176 us pulse length and 100 ms interval. Transfected parasites were recovered in D10F for roughly 24 hours, after which selection media was added.

### Ribosome Profiling and RNA Sequencing

ME49 parasites were mechanically released from host cells by scraping and syringe-releasing and used to infect 2 x 15-cm dishes of confluent human foreskin fibroblasts (HFFs) at an MOI of ∼2 with fresh D10F media. Parasites were harvested 48 hours post infection. For sample harvest, both 15 cm dishes per sample were placed on ice and washed twice with ice-cold PBS containing 100 ug/mL cycloheximide. 1 mL lysis buffer (20 mM HEPES pH 7.5, 100 mM KCl, 5 mM MgCl_2_, 1% Triton X-100, 2 mM DTT, 100 ug mL^-1^ cycloheximide, 500 U mL^-1^ RNaseIn Plus (Promega), and cOmplete Protease Inhibitor Cocktail (1 tablet per 10 mL buffer, Roche)) was added, and a cell lifter was used to dissociate HFFs with intracellular parasites from the dish, combining both dishes into the 1 mL sample. Lysates were passed through a 27-gauge needle 3 times and then a 30-gauge needle 3 times before incubating on ice for 5-10 minutes. Debris was pelleted by centrifugation at 1300xg and 4°C for 10 minutes, and supernatant was transferred to fresh tubes. The OD_260_ of each sample was measured, and samples were flash-frozen in liquid nitrogen and stored at −80°C. Two biological replicates were harvested.

For ribosome profiling, RNase I (Thermo Fisher) concentration was empirically tested, after which 40 U per 1 OD_260_ was used going forward. Sucrose gradient preparation, fractionation, collection of 80S ribosomes, and RNA fragment purification were performed as described in Xiang *et al*.^70^ For matched RNA-seq, probes for depletion of *Toxoplasma* rRNAs from total RNA samples were designed through NEB (**Supplementary Data 1**), and an equal ratio of human and *Toxoplasma* probes (2 uL each per reaction) were used for rRNA depletion with the NEBNext rRNA Depletion Kit (Human/Mouse/Rat, NEB, E6310L). Spike-in of GFP, Renilla luciferase, and Firefly luciferase mRNAs were added to the total RNA samples. Total RNA was fragmented with RNA Fragmentation Buffer (2 mM EDTA, 12 mM Na_2_CO_3_, 88 mM NaHCO_3_) by incubating at 95°C for 20 minutes, and fragmented RNA was purified by ethanol precipitation. Ribosome footprint and total RNA libraries were processed in parallel according to an established protocol.^71^ Briefly, 27 nt and 33 nt ^32^P-labelled size markers were used for size selection of RNA on a 10% urea gel. The 3′ end was dephosphorylated with T4 PNK (NEB) before ligation of the 3′ adenylated adapter (5′-App-NNNNCTGTCTCTTATACACATCTCCGAGCddC-3′). rRNA was depleted from ribosome footprint samples with a cocktail of biotinylated oligonucleotides complementary to human rRNAs (https://bartellab.wi.mit.edu/protocols.html) before another round of size selection. The 5′ end was phosphorylated using gamma-ATP and T4 PNK, followed by ligation of the 5′ adapter (5′-CCUACACGACGCUCUUCCGAUCUNNNN-3′) and a final size selection for ligated products. Libraries were reverse transcribed with SuperScriptIII (Invitrogen) and primer **P1** and amplified using Kapa HiFi polymerase (Roche) and barcoded primers (**P2-P9**). We performed two biological replicates. Sequencing runs were performed on either an Illumina NovaSeq (50×50 paired end cycles) or Illumina HiSeq (50 cycles).

### Read Processing and Genome Alignment

Reads from both sequencing runs were treated as single-end for compatibility. Adapters were trimmed with cutadapt (v3.7)^72^ and options -u 4 -a NNNNCTGTCTCTTATACACATCTCCGAGC --match-read-wildcards --minimum-length=15 and --nextseq-trim=20 or -q 20 for NovaSeq and HiSeq reads, respectively. Genomic and transcript fasta files and gff files for *T. gondii* ME49 were downloaded from ToxoDB Release 68 and the *H. sapiens* versions were downloaded from HostDB Release 68.^32^ STAR (v2.7.1a)^73^ was used to map reads to a metagenome of *T. gondii, H. sapiens*, size markers, and spike-in sequences with settings --outFilterType BySJout --outSAMattributes All --alignIntronMax 25000 --outSAMtype BAM SortedByCoordinate --outFilterMultimapNmax 1. Bigwig files were generated with bamCoverage^74^ and bin size of three nt (with a 12 nt offset for ribosome footprints) and visualized in IGV.^75^

### *Toxoplasma* 5′ UTR Annotation and Transcriptome Alignment

A custom *Toxoplasma* transcriptome annotation was generated to include representative 5′ UTR isoforms expressed in our data. For ToxoDB as well as Markus Tz TSS and Bz TSS transcript isoforms,^12^ 5′ UTRs were split into windows of three nt using bedtools^76^ makewindows, and total RNA coverage was calculated across these windows for each model using bedtools coverage -counts -s. We calculated cumulative read coverage and D for each model, where D is the ks.test statistic from comparison of a normalized, even distribution against read coverage for the model normalized to the UTR length and total cumulative sum. D values were compared for all models for a given gene, and the model with the lowest D value was selected as the representative isoform, except in the following instances: 1) in cases where the difference between D values was less than 0.05, we selected the model with the longest length; 2) in cases where all models had a D value greater than 0.4, the result was deemed inconclusive, and the 5′ UTR was excluded from further analyses. RPKM was calculated from counts generated with featureCounts, and transcripts with RPKM > 5 in both replicates were considered expressed. CDS and 3′ UTR annotations were used directly from ToxoDB models. In cases where there were multiple annotated CDS isoforms, the longest isoform was selected. A modified GFF was created with these gene models, and a spliced transcriptome fasta was created using gffread -w.

For the *H. sapiens* transcriptome annotation, canonical transcripts were identified from the UCSC genome browser^77^ knownCanonical option, and the *H. sapiens* transcriptome was filtered to include only the canonical transcript for each gene. The transcript with the longest CDS was selected for genes with multiple canonical transcripts or no canonical transcripts.

Genome-aligned reads were re-aligned to spliced *H. sapiens* and *T. gondii* transcriptomes with STAR options --outFilterMismatchNmax 2 --outFilterType BySJout --outSAMattributes All --outSAMtype BAM SortedByCoordinate.

### Translation Efficiency Analysis

Total mRNA and ribosome footprint reads mapping to the CDS of each transcript, excluding the first and last three codons, were counted with featureCounts^78^ and parameters GTF.featureType=“CDS”, useMetaFeatures=F, readShiftType = “downstream”, readShiftSize = 12, read2pos = 5, strandSpecific = 1. Ribosome footprint and mRNA RPKM were calculated for each gene using DESeq2,^79^ and translation efficiency (ribosome footprint RPKM / mRNA RPKM) was calculated for each biological replicate and then averaged.

### Construction and Testing of MPRA Vector

Cloning was performed using Q5 polymerase (NEB) and NEBuilder HiFi DNA Assembly Master Mix. To assemble pMLP140 (*pTub1-mNG-CDPK3 3*′ *UTR/pMIC2-mKate2-P2A-DHFR-DHFR 3*′ *UTR/Neutral Locus-targeting cassette*), the following segments were assembled: 1) neutral locus^52^ cassette (P10) was designed and synthesized based on the HiT vector strategy;^80^ 2) *pTub1-mNG* was amplified with primers P11/P12; 3) *CDPK3* 3′ UTR (GenBank:ON312869.1) was amplified with primers P13/P14; 4) *pMIC2*^81^ was amplified with primers P15/P16; 5) *mKate2* was amplified with primers P17/P18; 6) *P2A-DHFR*, plasmid backbone, and *U6* promoter was amplified from pALH459^82^ with primers P19/P20. To generate a digestible backbone for new 5′ UTR insertion, oligo P21 containing tandem BaeI restriction sites was cloned into the pMLP140 backbone amplified with P22/P23. To facilitate assessment of plasmid digestion, a stuffer fragment was amplified with primers P24/P25 and inserted between the BaeI sites, creating plasmid pMLP141. 5′ UTRs were amplified from parasite cDNA with primers introducing homology to *pTUB1* and *mNG* (P26-P35) and Gibson assembled into BaeI-digested plasmid pMLP141.

Plasmids were linearized by digestion with BsaI-HFv2 (NEB) and co-transfected with the pSS014 Cas9-expression plasmid (GenBank:OM640002) into RH parasites. Following recovery and then three passages under pyrimethamine selection (1.5 uM), populations were assessed by flow cytometry using a Miltenyi MACSQuant VYB. To test the effect of maintaining 50 nt of the *TUB1* 5′ UTR, we performed Gibson assembly of two fragments: 1) pMLP140 was amplified with primers P36/P37; 2) original plasmids with the *TGME49_207980* or *TGME49_211250* UTRs were amplified with primers P38 and P39 or P40, respectively. Transfections and flow cytometry analysis were performed as above. Plasmid pMLP160, which was ultimately used as the MRPA vector, was Gibson assembled by amplifying the BaeI cassette and fluorophore sequences from pMLP141 with primers P38/P41, and the extended *pTUBTUB1 5*′ *1/UTR* and remaining backbone with primers P37/P42.

### Library Assembly

A 5′ UTR library synthesized by Agilent (G7651A) was resuspended at 5 ng uL^-1^, amplified with iProof (Bio-Rad) and primers P43/P44 using 1 ng of template per 40 uL reaction, and gel extracted (Zymo). Vector pMLP160 was digested with BaeI (NEB) and gel extracted. The amplified library was Gibson assembled into BaeI-digested-pMLP160 (HiFi, NEB), dialyzed against water for 15 minutes, and electroporated into MegaX DH10B T1R electrocompetent cells (Invitrogen), which were grown on solid agar plates overnight. The library was harvested and frozen as pellets at −80C before maxiprepping (Zymo). The library was then linearized with BsaI-HF (NEB) and dialyzed against water. Plasmid pSS014 was maxiprepped.

### MPRA Screening

RH parasites were grown in 9 x 15-cm dishes with D3C media. 15 transfections were performed, with each transfection using 2×10^8^ parasites, 100 ug BsaI-digested library, and 50 ug Cas9 plasmid (pSS014) in 600 uL Cytomix. Transfections were pooled and split evenly across 15 x 15-cm dishes with D10F media. After 24 hours of recovery, the media was changed to D10F with 1.5 uM pyrimethamine and 10 ug mL^-1^ DNase I. At the first passage, any intact monolayer was scraped and parasites were pooled and concentrated before being evenly split across 15 x 15-cm dishes with fresh D10F + pyrimethamine media. At the second passage, parasites were passed into 10 x 15-cm dishes with 3×10^7^ parasites per plate and D10F + pyrimethamine media. At the third passage, parasites were passed onto 5 x 15-cm dishes with 4×10^7^ parasites per plate in D10F media. The remaining parasites were then harvested for FACS. Parasites were mechanically released by scraping and syringe-lysis and passed through a 5 uM filter. Aliquots of 1×10^8^ parasites were pelleted and frozen in liquid nitrogen for storage at −80°C until downstream processing alongside sorted samples. Parasites for sorting were resuspended in PBS + 2% IFS and sorted on a BD FACS Aria II. Sorted parasites were frozen, with at least ∼4-5 million parasites sorted into each bin. At the next lysis, sorting was repeated for a total of two biological replicates. DNA and RNA from both replicates were extracted with the DNA/RNA Allprep Mini kit (QIAGEN). For RNA, samples were treated with Turbo DNase (Ambion) and reverse-transcribed with Superscript III (Invitrogen) and an mNG-specific primer containing an 8-nucleotide unique molecular identifier (P45). RNA clean-up was performed with RNAClean XP beads (Beckman Coulter) before amplification with P46 and one barcoded primer (P47-P56). For DNA, samples were amplified with P57 and one barcoded primer (P58-P68). We also amplified the input plasmid library for reference. All amplified libraries were gel-extracted and sequenced on an Element Aviti 150×150 with custom primers for reads 1 and 2 and i7 indexing (P69-P71), and standard Illumina i5 index primer for sequencing of UMIs.

### MPRA Analysis

Reads were trimmed using cutadapt and paired-end reads were merged into one sequence with fastq-join. To include UTR sequences that were not in the designed library but may have resulted from library synthesis errors, all unique reads from any sample with length 170-190 nucleotides were identified. UTR sequences were used for downstream analysis if they were detected in both DNA replicates of unsorted parasites and the input plasmid library. Scripts adapted from Countess^83^ were used to count DNA or RNA reads (exact matches) and UMIs from RNA reads. Due to difficulty recovering high-quality RNA libraries from FACS samples, we analyzed the RNA from bulk populations only. To exclude sequences that may affect mNG expression at the level of transcription rather than translation, we used DESeq2 with standard parameters to compare both replicates of the de-duplicated bulk RNA read counts with bulk DNA read counts. To remove UTRs with disparate RNA and DNA levels, UTRs with adjusted *p* < 0.05 were excluded from further analysis. DNA read counts for the remaining UTRs were then used going forward to evaluate parasite distributions across bins. We required UTRs to have at least 25 reads in each bulk replicate, and a sum of at least 25 reads across all 4 sorted bins in each replicate. For each sample, reads were normalized to counts per million (cpm), and for each bin, UTRs were normalized to their corresponding bulk cpm. To calculate a representative MPRA score for each UTR, reads from both replicates were summed and a normal distribution was fit to the normalized read counts, with x-axis coordinates: mKate^+^/mNG^-^ = 0, bottom 15% mNG = 1, middle 15% mNG = 2, top 15% mNG = 3. The average value of this distribution (μ) is the MPRA score.

Construction of the random forest model was performed with scripts from Cao, Novoa, Zhang, *et al.*^50^ with the following modifications to features defined in the python script: 1) UTR codon frequency features were limited to open reading frames; 2) a Kozak similarity score feature was added; 3) descriptive uORF metrics including maximum and minimum uORF lengths, distance from CDS start to upstream features, and classification of uORF types (i.e. cuORF, ouORF, eUORF) were added; 4) additional nucleotide composition features were added to assess skew (denoted as “ratio”) between each pair of nucleotides; 5) 5′ cap folding energy was calculated using the first 100 nt of each UTR, including the constant *TUB1* region. We note that in the source code, the energies denoted as “Gquad” are calculated with standard RNAfold parameters and thus are redundant. The random forest model was constructed with ranger^84^ in R, using 100 trees during cross validation and 500 trees for the final model. We calculated Shapley values using fastshap^85^ with nsim=100. Scripts for the genetic algorithm were also adapted from Cao, Novoa, Zhang, *et al.*^50^ and used the GA package.^86^ We ran iterations of the GA optimizing for high or low TE with different starting sequences and parameters, including population size, probability of mutation, and probability of crossover. The highest or lowest TE sequence from each iteration was saved. These sequences were used as starting populations for another round of the GA. Representative sequences were selected from the highest and lowest TE results in both the first and second round of optimization.

### Cloning and Measurement of Synthetic 5′ UTR Expression

Synthetic UTRs with homology arms to mNG and the *TUB1* 5′ UTR were ordered as gBlocks from IDT (P72-P78) and cloned by Gibson assembly into BaeI-digested pMLP160. Plasmids were linearized by digestion with BsaI–HF and transfected into RH parasites. After recovery and several passages of growth under drug selection, The *pTUB1-5′ UTR-mNeonGreen-3′ UTR* locus was amplified from parasite gDNA with primers P79/P80, gel purified, and subjected to Sanger sequencing to confirm UTR identity.

Fluorescence intensity was measured after drug selection and across at least three passages with a Miltenyi MACSQuant VYB. For RT-qPCR analysis, intracellular parasites were scraped and pelleted before being flash-frozen in TRIzol for storage at −80°C. RNA was isolated by sequential TRIzol extraction, treated with Turbo DNase, and quantified by a NanoDrop spectrophotometer. First-strand cDNA synthesis was performed with random hexamer primers (Thermo Scientific) and SuperScriptIII (Invitrogen) using 2 ug of RNA. qPCR primers were assessed for efficiency (90%-110%) and specificity. qPCR was performed using 2X PowerUp SYBR Green Master Mix (Thermo Fisher) on a QuantStudio6 real-time PCR system (Applied Biosystems) using primers P81/P82 for mNG, P83/P84 for mKate2-P2A, and P85/P86 for ACT1, in standard cycling mode with T_m_ = 60℃.

### Data Availability

Sequencing data have been deposited in the Gene Expression Omnibus under accession numbers GEO:GSE302107 for ribosome profiling and GEO:GSE302108 for the MPRA.

## ACKNOWLEDGEMENTS

We thank Sumeet Gupta and the Whitehead Institute Genome Technology Core as well as the Flow Cytometry Core for key technical support; Jimmy Ly for consultation on ribosome profiling; Heather Keys for consultation on MPRA experimental design; and the Whitehead Bioinformatics and Research Computing Core, particularly George Bell, for consultation and bioinformatics support. We relied on VEuPathDB resources for this work and we thank all its contributors. A.S. was supported by a fellowship from the Helen Hay Whitney Foundation. This work was supported by grants AI144369 and AI158501 from NIH and 1021330 from Burroughs Wellcome Fund to S.L.

## DECLARATION OF INTERESTS

D.P.B. has equity in Alnylam Pharmaceuticals, where he is a co-founder and an advisor.

---

**Supplementary Figure 1.**
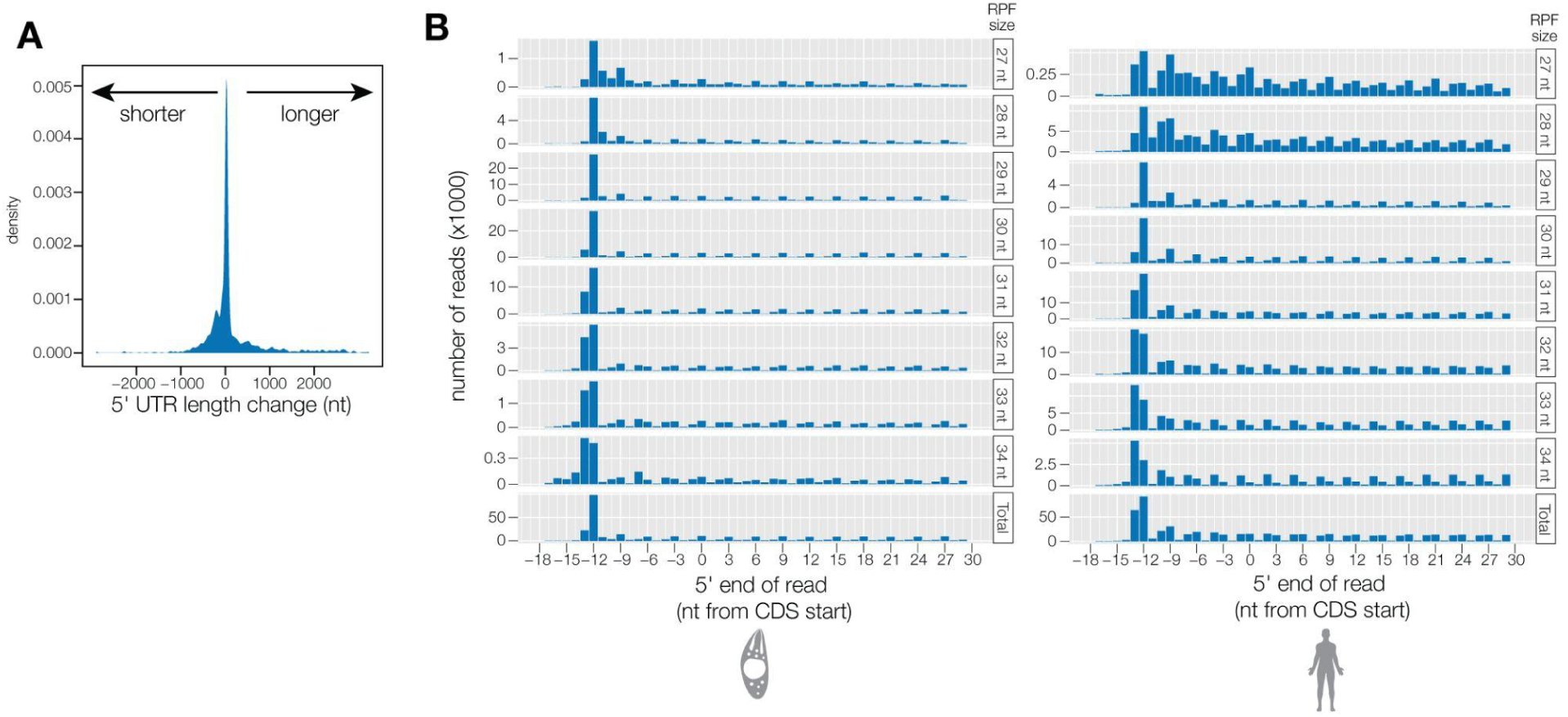
Changes to 5′ UTR models and assignment of ribosome P-sites. (**A**) Changes in 5′ UTR length (shown in nucleotides) following our re-annotation as compared to ToxoDB v68. (**B**) Ribosome footprints (RPFs) around the CDS start, grouped by length of RPF fragment. The most common peak for each RPF size was selected as the P-site offset for that footprint length.

**Supplementary Figure 2.**
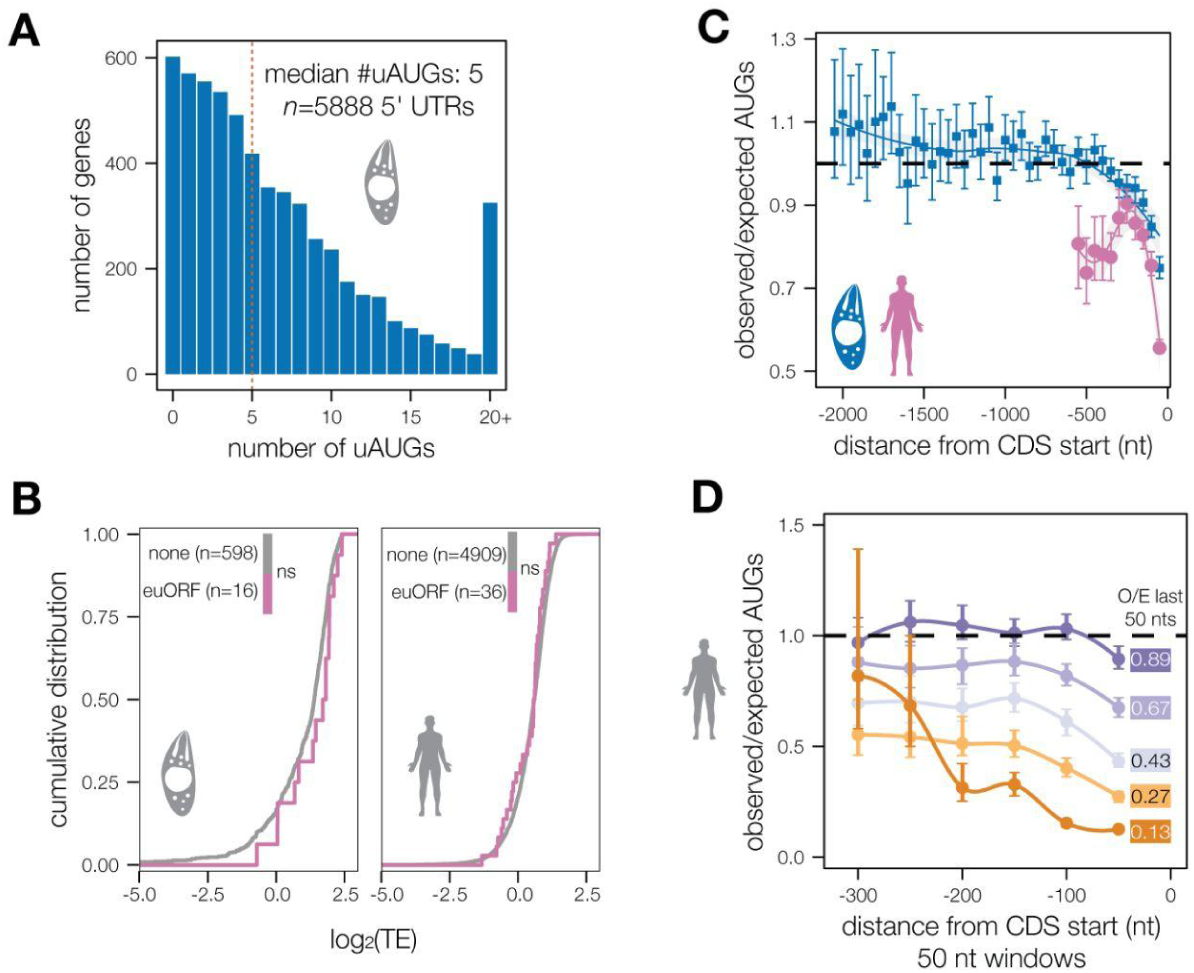
Additional analyses of uAUG distribution and effects on translation. (**A**) Distribution of the number of uAUGs in *Toxoplasma* 5′ UTRs with updated gene models. (**B**) Comparison of TE for transcripts with no uAUGs or one uAUG that is in-frame with the CDS start, forming an N-terminal extension (euORF). Wilcoxon rank sum test. (**C**) Observed/Expected uAUG ratio in 50 nucleotide (nt) windows for all *Toxoplasma* or human 5′ UTRs. (**D**) Observed/Expected uAUGs in human transcripts across TE quintiles, as in Figure 3G.

**Supplementary Figure 3.**
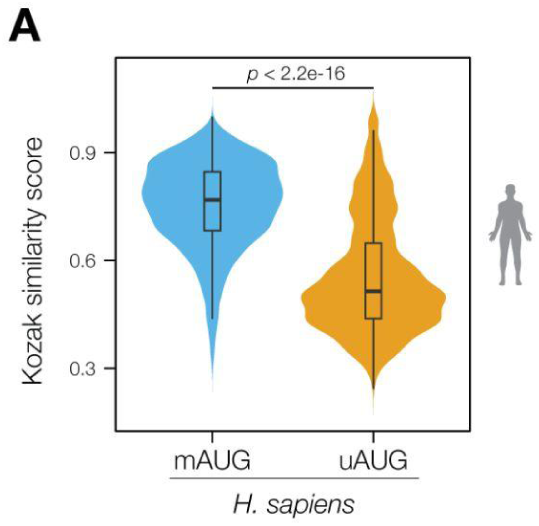
Human Kozak similarity scores. (**A**) Comparison of human Kozak similarity scores for mAUGs and uAUGs. Wilcoxon rank sum test.

**Supplementary Figure 4.**
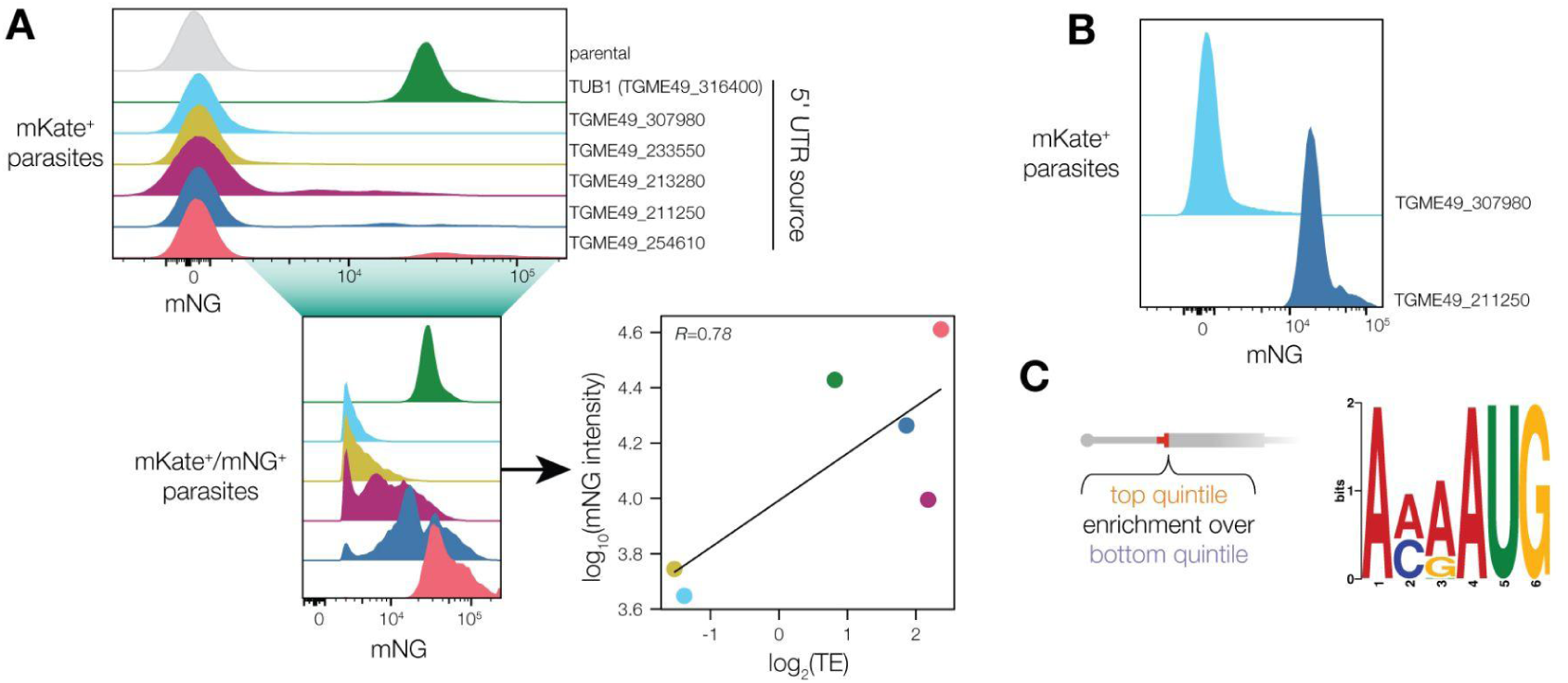
MPRA reporter development and analysis. (**A**) Development of reporter construct. Flow cytometry was used to measure mKate and mNG intensity. After gating for mKate-positive parasites in populations transfected with reporters containing 5′ UTRs inserted directly at the *TUB1* transcriptional start site (TSS), mNG expression was found to have a bimodal distribution. Inset shows the distribution of fluorescence intensity for mNG-positive parasites only. Scatter plot displays the correlation between the median mNG intensity of this mKate-positive/mNG-positive population and the endogenous TE for the corresponding UTR as measured by ribosome profiling. (**B**) mNG intensity by flow cytometry of mKate-positive parasites for reporters in which the 5′ UTR was inserted 50 nucleotides after the *TUB1* TSS, resolving mNG expression into a unimodal distribution. (**C**) STREME logo of the top motif identified in a discriminative search of sequences surrounding the CDS start (*E*-value: 1.3e-122). Top quintile MPRA score sequences were used as input, with bottom quintile MPRA score sequences as control.

**Supplementary Figure 5.**
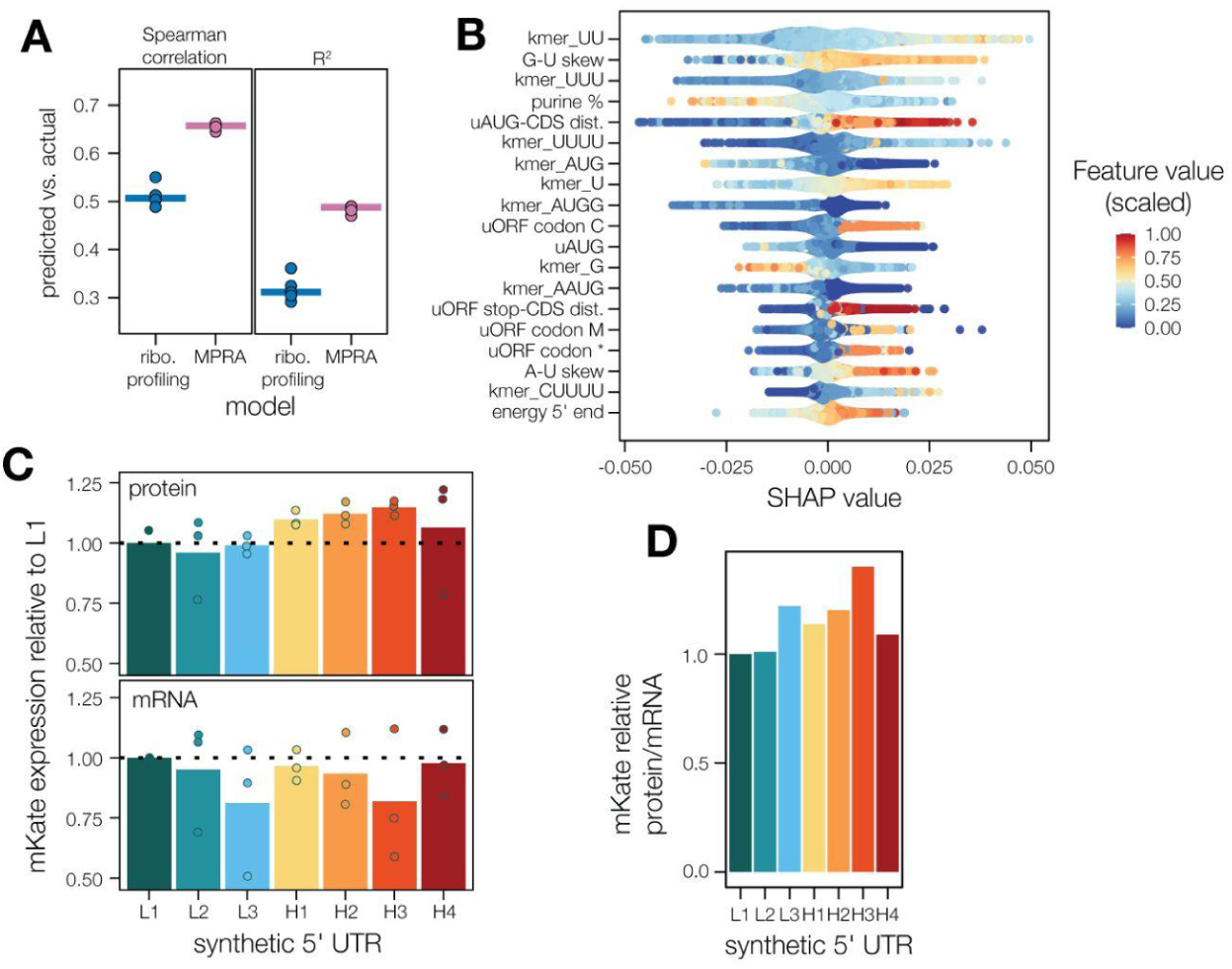
Model evaluation and controls for synthetic 5′ UTR reporters. (**A**) Comparison of performance of random forest models trained and tested on ribosome profiling data or MPRA data. Each model was subjected to stratified five-fold cross-validation in which the model is tested on unseen data. Individual points represent correlation metrics for each fold, and the median is represented by a bar. (**B**) Relationship between the values of top features and their Shapley values. (**C**) Quantification of mKate protein and mRNA levels as in Figure 6D. Raw C_t_ values are provided in **Supplementary Data 7**. (**C**) mKate pseudo-TE values as calculated in Figure 6E.

## Notes

http://www.ncbi.nlm.nih.gov/geo/query/acc.cgi?acc=GSE302107

http://www.ncbi.nlm.nih.gov/geo/query/acc.cgi?acc=GSE302108

